# Population structure limits parallel evolution

**DOI:** 10.1101/2021.01.26.428263

**Authors:** Bohao Fang, Petri Kemppainen, Paolo Momigliano, Juha Merilä

## Abstract

Population genetic theory predicts that small effective population sizes (*N_e_*) and restricted gene flow limit the potential for local adaptation. In particular, the probability of evolving similar phenotypes based on shared genetic mechanisms (i.e. parallel evolution), is expected to be reduced. We tested these predictions in a comparative genomic study of two ecologically similar and geographically co-distributed stickleback species *(viz. Gasterosteus aculeatus* and *Pungitius pungitius*). We found that *P. pungitius* harbours less genetic diversity and exhibits higher levels of genetic differentiation and isolation-by-distance than *G. aculeatus.* Conversely, *G. aculeatus* exhibits a stronger degree of genetic parallelism across freshwater populations than *P. pungitius:* 2996 *vs.* 379 SNPs located within 26 *vs* nine genomic regions show evidence of selection in multiple freshwater populations of *G. aculeatus* and *P. pungitius*, respectively. Most regions involved in parallel evolution in *G. aculeatus* showed increased levels of divergence, suggestive of selection on ancient haplotypes. In contrast, regions involved in freshwater adaptation in *P. pungitius* were younger, and often associated with reduced diversity. In accordance with theory, the results suggest that connectivity and genetic drift play crucial roles in determining the levels and geographic distribution of standing genetic variation, providing evidence that population subdivision limits local adaptation and therefore also the likelihood of parallel evolution.

## Introduction

Parallel evolution – defined here as the evolution of similar phenotypes in multiple independently colonised populations via selection on alleles that are identical by descent – is considered to be strong evidence for the role of natural selection in evolutionary change (Schluter et al. 2004). Adaptation from standing genetic variation (SGV) is thought to be the dominant route to parallel evolution among recently diverged populations (Conte et al. 2012; Ord & Summers 2015). Since genetic drift erodes SGV and barriers to gene flow can prevent beneficial alleles from reaching populations adapting to specific local habitats (Barrett & Schluter 2008; Feulner et al. 2013; Lenormand 2002), both the effective population size (*N_e_*) of the ancestral population (MacPherson & Nuismer 2017; Thompson et al. 2019) and gene flow (Bailey et al. 2017; Lee & Coop 2017; Ralph & Coop 2015) are expected to play key roles in determining the probability of parallel evolution. Both factors are expected to affect the heterogeneity in the geographic distribution of SGV across the distribution ranges of species. However, despite increasing interest to understand the drivers of parallel evolution over the past decade (e.g., Arendt & Reznick 2008; Barghi et al. 2019; Bolnick et al. 2018; Conte et al. 2012; Elmer et al. 2014; Elmer & Meyer 2011; Rosenblum et al. 2014; Stern 2013; Stuart et al. 2017), little effort has been placed to investigate the role of geographic heterogeneity in SGV (but see: Fang et al. 2020a; Kemppainen et al. 2021; Lee & Coop 2017). One way to test whether heterogeneity in pools of SGV determines the probability of parallel evolutionary responses is to investigate genetic structure and local adaptation in pairs of co-distributed and ecologically similar species that differ in their dispersal potential and population size, and hence, in the degree of heterogeneity of their pools of SGV.

The three-spined stickleback (*Gasterosteus aculeatus*) is an iconic model species used to study genetic parallelism. A multitude of studies has shown that the independent colonization of freshwater habitats across the global distribution range of this species has led to substantial, and often (but not always) parallel marine-freshwater associated genetic differentiation (Colosimo et al. 2005; DeFaveri et al. 2012; DeFaveri et al. 2011; Fang et al. 2020a; Hohenlohe et al. 2010; Hohenlohe & Magalhaes 2019; Jones et al. 2012). Despite the near circumpolar distribution of *Gasterosteus* sticklebacks, only three taxonomically valid species have been recognized in this genus (*viz*. *G. aculeatus, G. wheatlandi, G. japonicus*; Eschmeyer et al. 2017). In contrast, the circumpolarly distributed stickleback fishes in the genus *Pungitius* harbour at least eight taxonomically valid species (Eschmeyer et al. 2017; Guo et al. 2019; Takahashi et al. 2016), and there is evidence that the level of genetic differentiation among local populations in this genus greatly exceeds that seen among local populations of *Gasterosteus* (DeFaveri et al. 2012; Kemppainen et al. 2021; Merilä 2013; but see: Raeymaekers et al. 2017). Thus, comparative genetic studies of these two co-distributed species can provide an opportunity to gain novel insight into how differences in population structure, and thus in the distribution of SGV, may translate into differences in the probability of genetic parallelism. However, apart from two geographically restricted studies (DeFaveri et al. 2012; Raeymaekers et al. 2017), there has been no attempt to study and compare the levels of genetic variability and divergence among *Gasterosteus* and *Pungitius* taxa in a quantitative manner in a broad geographic context.

There is generally a greater abundance of three-than nine-spined sticklebacks in the sea (e.g., Cowen et al. 1991; Jurvelius et al. 1996; Ojaveer et al. 2003; Quinn & Light 1989) which is likely to limit gene flow and contribute to substantial genetic isolation by distance (IBD) in marine nine-spined sticklebacks. Consistent with this, earlier work suggests that the pool of SGV is indeed reduced and more fragmented in nine-compared to three-spined sticklebacks (DeFaveri et al. 2012; Kemppainen et al. 2021; Merilä 2014). To gain a holistic view of factors that influence such differences in SGV, using comprehensive geographic sampling and high-density population genomic data, we first re-assess the differences in both phylogenetic histories and population demographic parameters between these two species. We then formulate and test the hypothesis that due to a higher geographic heterogeneity in SGV in nine-spined sticklebacks, this species will show a lower prevalence of parallel evolution in response to freshwater colonisation than the three-spined stickleback.

## Materials and methods

### The study species

The two study species are ecologically very similar and are frequently syntopic in both marine and freshwater habitats (e.g., Copp & Kovac 2003; DeFaveri et al. 2012; Ojaveer et al. 2003; Raeymaekers et al. 2017). However, there is a tendency for the three-spined stickleback to be more common in marine habitats, and for the nine-spined stickleback to be more common in freshwater habitats (Wootton 1976; Wootton 1984). Both species are small (typically < 100 mm), with similar lifespans (Baker 1994; DeFaveri & Merilä 2013; DeFaveri et al. 2014) and breeding habits (Wootton 1976; Wootton 1984), and exhibit male parental care; males build and attend nests in which multiple females can lay their eggs (Wootton 1976; Wootton 1984). Females of both species can lay single or multiple clutches of *ca.* 100-500 eggs per breeding season (Baker 1994; Heins & Baker 2003; Herczeg et al. 2010; Wootton 1976; Wootton 1984). Hence, we do not expect a large variation in levels of SGV between the two species due to differences in life-history traits (cf. Ellegren & Galtier 2016; Romiguier et al. 2014).

### Sample collection and sequencing

The data set is composed of 166 (47 marine and 119 freshwater) three-spined stickleback and 181 (48 marine and 133 freshwater) nine-spined stickleback individuals. The 166 three-spined stickleback samples were the same as used in Fang et al. 2020a). Data for 75 nine-spined stickleback samples were retrieved from an earlier Restriction Site Associated DNA (RAD) sequencing study (Guo et al. 2019) and a whole-genome re-sequencing (WGRS) study (Feng et al. 2020). New samples of nine-spined sticklebacks were sequenced specifically for this study, including 23 samples using RAD sequencing (protocol following Guo et al. 2019, using the PstI enzyme) and 83 samples using WGRS at 10X coverage (protocol following Feng et al. 2020). In total, 63 populations (26 marine and 37 freshwater; sample sizes: 1–10) of three-spined stickleback and 36 populations (7 marine and 29 freshwater populations; sample sizes: 1–20) of nine-spined stickleback were included in this study. The sampling sites are shown in Fig. 1a and detailed sample information (population acronyms, sample sizes, lineages, sampling site coordinates, sequencing information etc.) are given in Supplementary Table 1.

**Figure 1.**
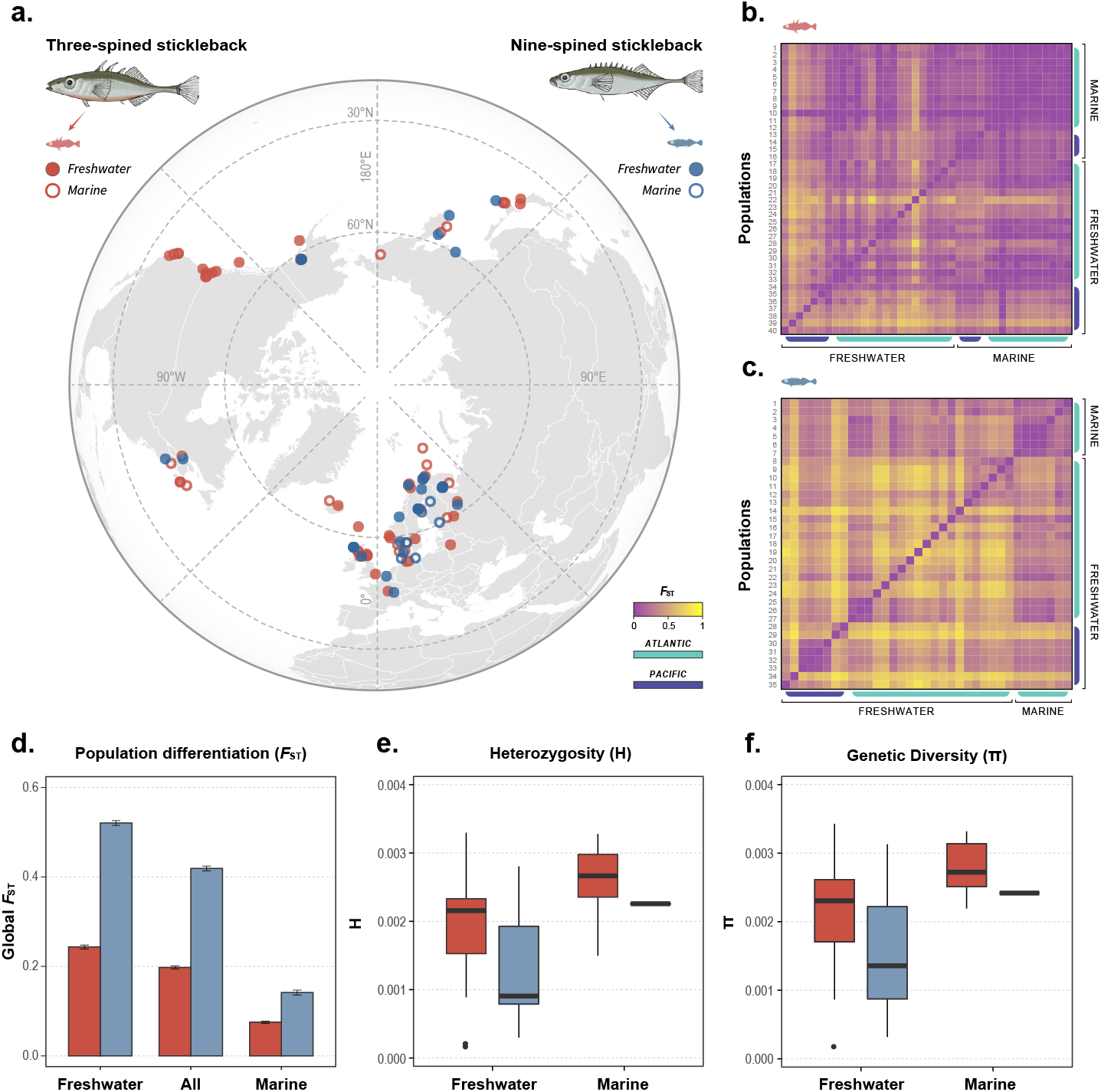
Three-spined sticklebacks exhibit higher levels of genetic variation and lower degree of population differentiation than nine-spined sticklebacks. (a) Map showing the species-specific sampling locations. (b, c) Pairwise population differentiation (*F*_ST_) of three- and nine-spined sticklebacks, respectively. Populations are specified in Supplementary Table 2. (d) Global *F*_ST_ of the two species. Error bars indicate 95% bootstrap confident intervals. (e, f) Boxplots of Heterozygosity *(H)* and nucleotide diversity *(π).* Admixed populations in nine-spined stickleback are excluded in (e-f). See Supplementary Fig. 2 for the estimates of Watterson’s theta *(θ)* and the effect of admixed populations on measures of genetic diversity. Generalized linear mixed model (GLMM) revealed significant differences in genetic diversity (e,f) between species and ecotypes (see Results).

According to the known phylogenetic relationships within each species, the three-spined stickleback samples were assigned to seven lineages: Eastern Pacific (EP), Western Pacific (WP), Western Atlantic (WA), White and Barents Seas (WB), North Sea & British Isles (NS), Baltic Sea (BS) and Norwegian Sea (NOR) (Fang et al. 2018); and the nine-spined stickleback samples were assigned to six lineages; Western Europe (WL), Eastern Europe (EL), Baltic & North Seas (BN; this lineage has been formed through admixture between the Western European [WL] and Eastern European [EL] lineages; Feng et al. 2020; Guo et al. 2019; Teacher et al. 2011), Far East (FE), North America (NA) and Alaska (ALA).

The new stickleback samples were collected during the local breeding seasons with seine nets and minnow traps. After euthanizing the fish with an overdose of MS-222, whole fish or fin clips were preserved in ethanol for DNA-extractions using the salting-out method (Sunnucks & Hales 1996). The sample collection in Finland was conducted with personal fishing licences and permissions from the landowners according to the Finnish Fishing Law (5§ 27.5.2011/600). In other countries, the sampling was performed under respective national licenses granted to the sample-providers. The study does not involve animal experiments according to the Finnish National Animal Experiment Board (#STH379A and #STH223A). For the newly sequenced nine-spined stickleback samples, the RAD-sequencing data and the WGRS data were obtained with the protocols given in Guo et al. 2019) and Feng et al. 2020), respectively.

### Genotype likelihood estimation

The same bioinformatics pipelines were applied to both species. For each species, all RAD and WGRS sequences were mapped to their respective reference genomes with BWA mem v.0.7.17 (Li & Durbin 2010). The reference genome of the three-spined stickleback was retrieved from the Ensembl database (release-92; Yates et al. 2020) and that of the nine-spined stickleback from (Varadharajan et al. 2019; ver. 6). Genotype likelihoods were estimated from the mapped reads using ANGSD v.0.93 (Korneliussen et al. 2014) with the same parameter settings for both species. Quality filtering parameters are explained in detail in the Supplementary Method 1. The raw output of genotype likelihoods from the 166 three-spined sticklebacks comprised 2,511,922 SNPs and those of the 181 nine-spined sticklebacks 7,938,802 SNPs. The difference in SNP numbers between species partly reflects the larger proportion of WGS samples in the latter species (80.1%) than in the former species (22.9%).

### Genetic diversity and differentiation

Genetic diversity within populations was estimated by computing population nucleotide diversity (*π*, Nei & Li 1979) and Watterson’s theta (*θ =4Neμ*, where *N_e_* is effective population size and *μ* the mutation rate, Watterson 1975) as well as individual heterozygosity (*H*, the proportion of heterozygous sites within an individual genome) with ANGSD and custom R-scripts (Supplementary Method 2). Since some of the sampled populations are known to be admixed (Feng et al. 2020; Guo et al. 2019; Supplementary Method 2), their genetic diversity was expected to be elevated. We report the results both when excluding and including the admixed nine-spined stickleback populations. In each species, the allelic differentiation *F*_ST_ (Weir & Cockerham 1984) was calculated over all samples, within marine and within freshwater ecotypes (global *F*_ST_ over all loci), and between populations (pairwise *F*_ST_). To do so, we used a subset of high-quality genotypes to estimate global *F*_ST_ for each ecotype and the pairwise *F*_ST_ between all populations with the R packages *hierfstat* (Goudet 2005) and *StAMPP* (Pembleton et al. 2013), respectively. Details of the methods used to estimate genetic diversity and differentiation are specified in the Supplementary Method 2.

The levels of genetic diversity (*H*, *π* and *θ*) between the two species were compared by fitting generalized linear mixed-effects models (GLMMs) in R using the packages *lme4* (Bates et al. 2014) and *Car* (Fox et al. 2012). The models treat species, habitat (Freshwater, Marine) and their interaction as fixed effects. The geographic region was set as a random effect to account for non-independence between populations across regions. Non-significant interactions were deleted from the final models. To test differences in global *F*_ST_ between species and habitats, we performed bootstrapping based on 10,000 resampled datasets in which each resample consisted of 1/3 of the markers and 1/3 of the samples to obtain the 95% confidence intervals.

### Isolation-by-distance

Tests for isolation-by-distance (IBD) were performed by regressing pairwise genetic distances (linearized *F*_ST_ = *F*_ST_ /[1 – *F*_ST_]; Slatkin 1995) against pairwise geographic distances between populations. Our sampling of the nine-spined sticklebacks from the Eastern Pacific region was very thin: only two freshwater and no marine populations were sampled. Therefore, to characterize IBD for different ecotypes (marine and freshwater populations), we performed the IBD tests on the European populations (see Supplementary Fig. 1d for sampling map), where both ecotypes for both species were available. Geographic distances were measured between marine populations based on the pairwise least-cost geographic distances across marine environments using the R Package *Marmap* (Pante & Simon-Bouhet 2013), and between freshwater populations based on world geodetic system with the R package *raster* (Hijmans & van Etten 2014). To test if the IBD relationships differed between the two stickleback species and habitats, the IBD regressions were fitted with maximum-likelihood population effects (MLPE) models to account for the non-independence of pairwise distances (Clarke et al. 2002) and slopes of the IBD regressions for both species were compared. The MLPE analyses were performed using the R packages *corMLPE* (Clarke et al. 2002) and *nlme* (Pinheiro et al. 2017).

The White Sea marine population of the nine-spined stickleback (RUS-LEV) has a close phylogenetic relationship with the marine populations of the Baltic Sea, since the latter originated from a post-glacial invasion from the White Sea over an area today occupied by land (Guo et al. 2019; Shikano et al. 2010a)this study [Fig. 3]). Therefore, there is a clear rationale to expect RUS-LEV to be an outlier in IBD analysis. We thus performed IBD analyses excluding the population RUS-LEV but we also report the results when including it in Supplementary Information 1.

**Figure 3.**
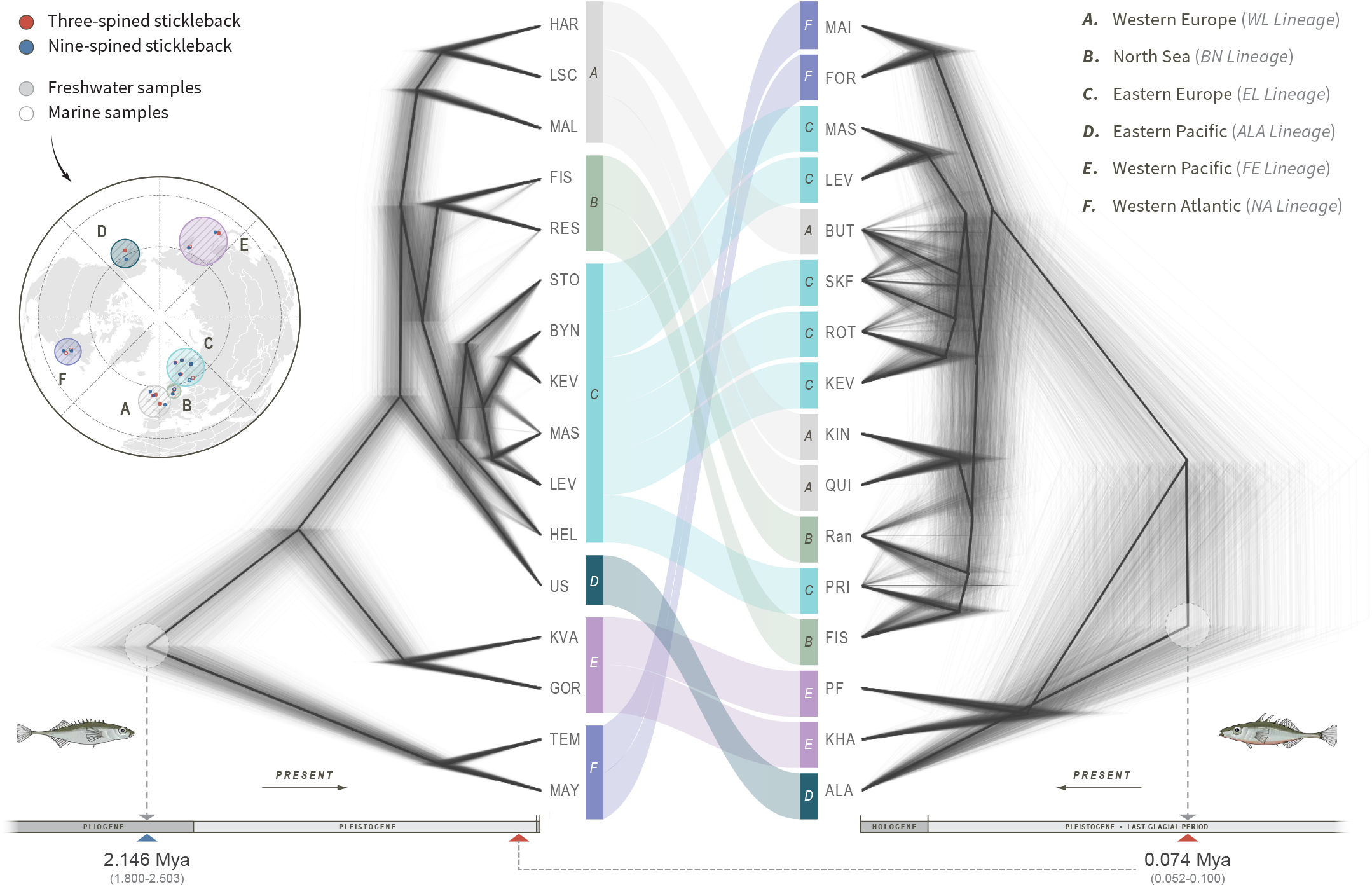
Time-calibrated phylogenies of three- and nine-spined sticklebacks inferred with SNAPP. The topology and divergence times of phylogenetic trees of nine-(left) and three-spined sticklebacks (right) are presented and compared using populations across the same, or geographically close, sampling sites. Colours correspond to the different lineages (A-F) of nine-spined sticklebacks. Arrows in the bottom indicate directions of timeline from past to present. The TMRCA (time to the most recent common ancestor) of three-spined sticklebacks is marked in the timeline of the nine-spined sticklebacks for comparison. Population identifiers were simplified for clarity; see Supplementary Fig. 1e for a map of sampled populations. The maximum-clade-credibility summary trees of the SNAPP phylogenies indicating divergence times and calibration points are given in the Supplementary Fig. 3.

### Comparative phylogenomic analyses

There is evidence to suggest that the probability of genetic parallelism decreases with increasing divergence time between taxa (Conte et al. 2012; Ord & Summers 2015). To assess differences in divergence times among populations of the two species, time-calibrated phylogenetic trees were constructed using genome-wide SNPs based on the multispecies coalescent model with the program SNAPP (Bouckaert & Bryant 2012; Chifman & Kubatko 2014).

For these analyses, we selected 16 paired sampling locations from where samples of both species were available, representing all major biogeographic regions within the two species’ distribution ranges (Fig. 3). SNAPP analyses were performed using filtered datasets of 12,022 SNPs for three- and 13,079 SNPs for nine-spined sticklebacks (bi-allelic SNPs > 10 kb apart, with no missing data, and posterior probability > 0.95%), following the protocols of Stange et al. 2018 and Fang et al. 2020b. The time calibrations were conducted using the divergence time estimates derived from Guo et al. 2019 and Fang et al. 2020b for nine- and three-spined sticklebacks, respectively. Detailed methods of SNP filtering and phylogenetic analyses are specified in the Supplementary Method 3.

### Comparative analyses of genetic parallelism

The degree of genetic parallelism in response to freshwater colonisation in three- and nine-spined sticklebacks was assessed in two steps. First, Linkage Disequilibrium Network Analyses (LDna; Fang et al. 2020a; Kemppainen et al. 2015; Li et al. 2018) was used to partition data into correlated sets of loci (LD-clusters), followed by linear mixed models (LMM) testing for associations between PC-coordinates based on loci from the LD-clusters and ecotype (treated as a binary trait) while controlling for *p*-value inflation due to relatedness and other confounding factors. For these analyses, only samples from the Atlantic region were used, and the data sets were normalised such that the same number of polymorphic loci were analysed for both species after accounting for differences in sequencing coverage and genetic diversity (see Supplementary Method 4). Ultimately, 882,125 and 1,355,325 SNPs (in the form of genotype likelihoods) were used in the downstream comparative LDna analyses in three- and nine-spined sticklebacks, respectively.

#### Complexity reduction using LDna analyses

Since LDna relies on pairwise LD-estimates among all loci, it is not feasible to consider all pairwise comparisons at once for large data sets. Instead, we adopted a nested approach, starting with LDna within windows within chromosomes (LDna-1, *sensu* Li et al. 2018; Supplementary Method 4), followed by LDna within chromosomes (LDna-2) and finally LDna for genome wide SNPs (LDna-3) as described in Fang et al. 2020a, with some modifications (an overview of this approach is given in Supplementary Fig. 2 and further details are given in Supplementary Method 4). First, LD-clusters were defined by a single parameter, the minimum number of edges in the cluster (|*E*|*_min_*), rendering the previously used *λ_lim_* parameter (that determines how different LD-signals are between different clusters) obsolete.

Second, instead of using one SNP (rSNP) to represent a cluster in the subsequent LDna-step, the final LDna-3 was based on a correlation matrix of *r*^2^ between PC1-coordinates (based on genotype likelihoods from all loci from a given LD-cluster; PCAngsd) (Meisner & Albrechtsen 2018) between all pairs of LD-clusters entering LDna-3 analyses. The first PC typically explains >>90% of the variation in each LD-cluster and in Supporting Information 5 we demonstrate that these coordinates can be regarded as “synthetic multi-locus alleles” (SMLAs). Third, all LDna-1 clusters with more than *SNP_min_* number of loci and that were not part of any LDna-2 cluster were also included in the final LDna-3 analyses. Fourth, LDna-3 clusters were determined by the parameter *Cor_th_*, which specifies the weakest link (*r_2_*-value) in a cluster that is allowed between LDna-1 and LDna-2 clusters entering LDna-3. As such, decreasing parameter values for |*E*|*_min_*, *SNP_min_* and C*or_th_* lead to many smaller clusters with few but highly correlated loci and vice versa.

There is a trade-off between *i*) “under clustering” (i.e. analysing many clusters that in reality reflect the same evolutionary phenomena leading to overly conservative corrections for multiple testing), and *ii*) “over clustering” (i.e. analysing fewer and larger clusters but each with sets of less correlated loci). While the latter leads to less conservative corrections for multiple testing (higher power), it likely also leads to weaker associations between the SMLAs and ecotype. Importantly, the parameter settings that maximises the power to detect a particular genomic region of interest will depend on both the data set (numbers of loci and the underlying LD-structure) and the genomic region in question. Here we solved this problem by testing a range of parameter settings for |*E*|*_min_* [10,20,40], *SNP_min_* [10,20,40] and *Cor_th_* [0.8,0.7,0.6,0.5] for both data sets. All LD-values were estimated by *ngsLD* (Fox et al. 2019) based on genotype likelihoods. Further details are given in Supplementary Method 4 and below.

#### Linear Mixed model analysis for testing associations between LD-clusters and ecotype

We used linear mixed models (LMM) to test for associations between ecotype and the genetic variation explained by loci in LD-clusters by regressing the SMLAs against ecotype treated as a binary trait [0,1]. Using LMM to test associations between genotype and phenotype has previously been shown to be analogous to using permutation to test for allele frequency difference between two groups (Kemppainen et al. 2017), with the major benefit of LMM’s being computational speed and the ability to account for confounding factors such as relatedness. When not accounting for any potential confounding factors, the LMM-approach used here produces test statistics that are highly correlated with both *FST* and the cluster separation score (CSS; Supporting Method 5; Supplementary Fig. 3), two commonly used metrics to detect genomic regions associated with parallel evolution in three-spined sticklebacks (Fang et al. 2020a; Jones et al. 2012; Kingman et al. 2020). We used a modified version of the restricted maximum likelihood (REML)-based method EMMAX (Efficient Mixed-Model Association eXpedited; Kang et al. 2010; Li et al. 2018) that allowed us to test for associations between SMLAs (rather than a single bi-allelic SNP at a time) from LD-clusters and ecotype (Supporting Method 6).

Two approaches to control for multiple testing were used: permutation (Li et al. 2018) (Supplementary Method 7) and the “*HS*” method aka false discovery rate “*fdr*” (Benjamini & Hochberg 1995). In addition, two methods to control for *p*-value inflation caused by relatedness were used: including a relatedness matrix ***A*** as a random effect (Kang et al. 2010) and genomic control (Price et al. 2010). Whenever *fdr* was used to control for multiplicity, we also iteratively estimated *p*-value inflation as the linear slope λ between observed and expected (under the null-hypothesis) −log10(*P*) values before and after removing significant LD-clusters from the data (the medians from all orthogonal combinations of *|E|_min_*, *SNP_min_* and *Cor_th_* were used). The reason for this was the exceptionally high proportion of the genome involved in parallel marine-freshwater differentiation that would have led to an overestimation of *λ*, especially in the three-spined stickleback (iterations stopped when no more or no less significant genomic regions were found compared to the previous iteration). Whenever *p*-value inflation was present (*λ*>1) all observed −log10(*P*) were divided by *λ* (prior to *fdr*), thus ensuring that no residual *p*-value inflation would exist in the data, also known as genomic control (GC; Price et al. 2010). For instance, *λ*=2 (high *p*-value inflation) means that a test with −log10(*P*)=10^−2^ after GC-correction is no longer significant (10^−2^/*λ*=0.1). Note that GC was seldom necessary when relatedness was accounted for and GC is not possible when permutation is used to control for multiplicity (Li et al. 2018).

All association analyses were corrected for *p*-value inflation and multiplicity using four approaches, *i*) including relatedness as a random effect and using *fdr* (“A+fdr”), *ii*) including relatedness as a random effect and controlling for multiplicity by permutation (“A+perm”), *iii*) ignoring relatedness and instead using GC to control *p*-value inflation, followed by *fdr* (“GC+fdr”) and *iv*) not including relatedness as a random effect but controlling for multiplicity using permutation (“perm”; with no possibility for GC).

Due to the two highly divergent lineages of nine-spined sticklebacks in the Atlantic (with some individuals being a result of admixture between them), we included lineage (WL, EL or ADMIXED) as a co-factor in the analyses for this species. This greatly reduced initial *p*-value inflation that otherwise would have caused many false positives or significantly reduced power to detect true significant associations following GC.

#### Assessing sensitivity to of association analyses to parameter settings

With three parameters for defining LD-clusters (|*E*|*_min_*, *SNP_min_* and *Cor_th_*) and four methods to correct for multiplicity and *p*-value inflation, the three- and nine-spined stickleback data sets were subjected to a total of 180 tests each. It is important to note, however, that all tests are applied to exactly the same data sets (i.e. tests within species are per definition not independent), such that the cumulative number of significant genomic regions found is quickly expected to reach an asymptote as more parameter combinations are tested, as shown in Supplementary Figure 4.

A genomic region was considered significant when at least ten unique loci from clusters significant at α=0.05 (after corrections) were also physically clustered in the genome as determined by single linkage clustering with a distance threshold of 500kb. All significant clusters from any of the 180 parameter/correction method combinations were included but most loci were found in LD-clusters in multiple such combinations. Based on this, we calculated a consistency score *C* for each putative outlier region, denoting the proportion of tests where a given genomic region was found significant, with *C*=1 indicating that a given region was significant in all 180 tests. Conversely, low C-scores are expected for outlier regions that are only detected in a restricted set of parameter/correction combinations. Note, however, that regions with low *C*-scores do not necessarily imply small effect sizes, although they can be correlated. Our wide range of parameter combinations was necessary to minimise the dependence between parameter values and the number of outlier regions detected in three- and nine-spined sticklebacks, despite vastly different levels of population structuring and potentially fundamentally different mechanisms underlying marine-freshwater parallelism. We deemed outlier regions with *C*<0.05 to be too sensitive to parameter settings to be considered further in downstream analyses (Supplementary Fig. 4).

#### Regional parallelism

We also performed EMMAX analyses separately for the geographic regions with large sample sizes of freshwater individuals for both species: Baltic Sea (18,13), North Sea (20,23), Norwegian Sea (21,23) and White & Barents Seas (31,38), with numbers in brackets indicating sample sizes for three- and nine-spined sticklebacks, respectively. The corresponding marine samples sizes were more variable for both species and were lacking altogether from the Norwegian Sea region for the nine-spined sticklebacks. However, for any genomic region associated with marine-freshwater parallelism, the expectation is that freshwater adapted alleles/haplotypes are found in high frequency in the freshwater populations where they are locally adapted. Conversely, in marine populations, we expect the low frequency of these alleles/haplotypes, regardless of geographic location (in contrast to neutral loci). Thus, in order to analyse and compare regional parallelism for both species fairly, we pooled all marine samples and contrasted them against freshwater samples from one of the four geographic regions at a time. These analyses were performed on SMLAs based on all unique loci from significant LD-clusters that mapped to each significant genomic region identified above. Since the power to detect significant associations depends on the data set, we only compared correlation coefficients as proxies for effect sizes, both when assuming all individuals are unrelated *(cor_unrl_*) and when including ***A*** as a random effect *(cor_unrl_*). This allowed us to assess which geographic regions contributed, and how much, for the overall marine-freshwater differentiation for each significant genomic region.

#### Divergence times and parallel evolution

Since genetic parallelism is expected to be a negative function of time since divergence (e.g., Conte et al. 2012), we further explored the correlation between divergence time and the level of genetic parallelism using seven freshwater population pairs from Europe (shown in Supplementary Fig. 5). In each species, we first extracted the divergence times (in Mya) between pairwise intra-specific populations based on the maximum-clade-credibility summary tree, using the R-package *ape* (Paradis et al. 2004). The level of genetic parallelism for each pair of freshwater-freshwater populations was estimated by counting the proportion (relative to the entire data set) of marine-freshwater associated LD-cluster (-log_10_(*p*) > 2) loci that grouped the two freshwater populations in the PCA (individuals from both freshwater populations were found in the in-group) for a given LD-cluster. Finally, the matrixes of pairwise intra-specific divergence times and levels of genetic parallelism were fitted using MLPE model similar to the IBD analyses (described above).

Since processes that govern diversity levels within genomes (background selection, mutation rate and recombination rate variation) are conserved between closely related populations (and species), different measures of diversity are correlated across the genomes of closely related populations (Dutoit et al. 2017). Here we take advantage of this correlation to detect whether outlier genomic regions have more ancient origins than neutral regions by testing whether genomic regions under selection show excess absolute divergence (*d_XY_*, Nei 1987) relative to the rest of the genome (*Δd_XY_*) as detailed in Supplementary Method 8.

## Results

### Genetic variation within populations

There were significant differences in levels of genetic diversity between the two species and habitats. Average heterozygosity (*H*) was significantly higher in three-spined than in nine-spined sticklebacks (GLMM: F_1,258.85_=91.33, *P*<0.001; Fig. 1e). Marine populations harboured higher heterozygosity than freshwater populations in both species (GLMM: F_1,257.14_=25.70, *P*<0.001; Fig. 1e). Both *π* and *θ* were also higher in the three-than in the nine-spined stickleback populations (*π*: GLMM, F_1,58.91_=10.34, *P*=0.002, Fig. 1f; *θ*: GLMM, F_1,58.98_=12.48, *P*<0.001, Supplementary Fig. 6a), and higher in marine populations than in freshwater populations (*π*: GLMM, F_1,58.98_=12.49, *P*<0.001, Fig. 1f,g; *θ*: GLMM, F_1,58.61_=7.25, *P*<0.01, Supplementary Fig. 6a). Species*habitat interactions were not significant in any of the analyses.

When incorporating nine-spined stickleback populations showing strong signatures of admixture (see Materials and methods) in the analyses, the differences in genetic diversity (*H*, *π* and *θ*) between habitats were still significant (*H*: GLMM: F_1,320.45_=54.59, *P*<0.001; *π*: GLMM, F_1,70.09_=9.24, *P*=0.003; *θ*: GLMM, F_1,69.93_=8.21, *P*=0.005; Supplementary Fig. 6b-d), but those between species were no longer significant (Supplementary Fig. 6b-d). This indicates that admixture has had a significant positive effect on genetic diversity in the admixed nine-spined stickleback populations. Indeed, admixed populations have significantly higher heterozygosity than non-admixed populations (GLMM, F_1,320.02_=75.82, *P* < 0.001; Supplementary Fig. 6e).

### Genetic differentiation among populations

The degree of genetic differentiation among nine-spined stickleback populations was significantly higher (global *F*_ST_ = 0.419, 95% CI: 0.414-0.424; Fig. 1d) than that of three-spined stickleback populations (global *F*_ST_ = 0.198; 95% CI: 0.194-0.201; Fig. 1d). This was true also when only considering populations from the same genetic clades in nine-spined sticklebacks (Supplementary Fig. 7). In both species, there was less differentiation among marine than freshwater populations (Fig. 1d). Furthermore, in the case of the nine-spined stickleback, IBD was significant in both marine and freshwater environments (MLPE, p≤0.01; Fig. 2). In the three-spined stickleback, IBD was significant in marine (MLPE, *P*<0.001; Fig. 2) but not in freshwater habitat (MLPE, *P*<0.26; Fig. 2). A comparison of the IBD slopes (MLPE regression coefficient *β*) revealed that the IBD in nine-spined sticklebacks was 23.9 times stronger in the marine habitat (MLPE, *β* = 1.2e-4 *vs.* 5.1e-6; Fig. 2).

**Figure 2.**
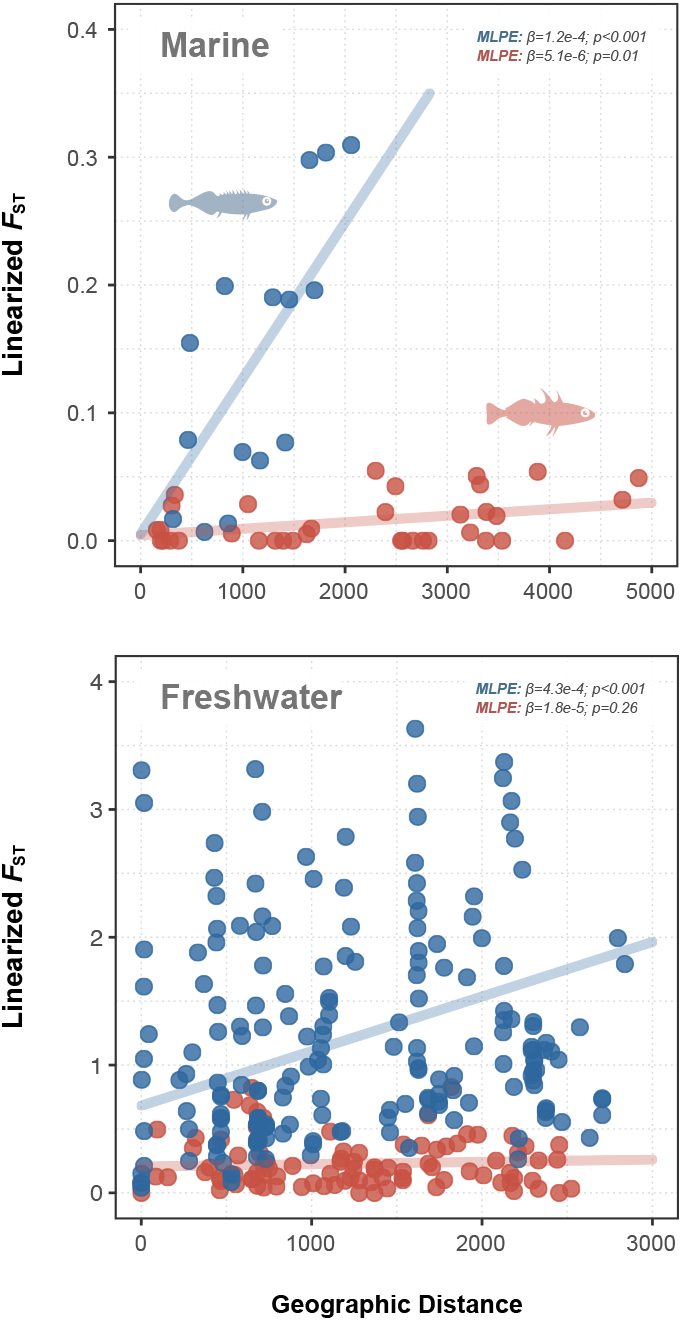
Isolation by distance (IBD). IBD was tested across (a) marine and (b) freshwater three-(red) and nine-spined stickleback (blue) populations using maximum-likelihood population-effects (MLPE) model. The slope of the regression coefficients *(β*) suggest stronger IBD in nine-than in three-spined sticklebacks. IBD comparisons were restricted to European populations (see Materials and Methods).

### Phylogenetic histories

The comparison of the time-calibrated phylogenies between three- and nine-spined sticklebacks (Fig. 3) revealed contrasting phylogenetic relationships and colonisation histories across their global distribution. The TMRCA of all lineages of nine-spined stickleback was 2.146 Mya in late Pliocene (95% HPD interval [hereafter in parenthesis]: 1.800-2.503 Mya), which is much older than that of the three-spined stickleback 0.074 Mya in late Pleistocene (0.052-0.100 Mya).

The most ancient lineage of nine-spined sticklebacks was from the Western Atlantic (F in Fig. 3). In contrast, the Western Atlantic clade of three-spined sticklebacks was among the youngest lineages, in line with earlier findings (Fang et al. 2018, 2020b). The most ancestral lineage in three-spined sticklebacks was from the Eastern Pacific clade (D in Fig. 3), whereas nine-spined sticklebacks from this area were more recently diverged (ALA lineage) with a divergence time close to the TMRCA of its European lineages (0.766 Mya [0.644-0.887 Mya]; Fig. 3, Supplementary Fig. 5).

The European three-spined stickleback populations have diverged recently (A, B and C in Fig. 3; 0.026 Mya [0.018-0.035 Mya]), with significant incomplete lineage sorting among them. In contrast, the European nine-spined stickleback populations had deep and clear lineage separation (three lineages [A, B and C] diverged 0.762 Mya [0.638-0.882 Mya]), with evidence for introgression between the Eastern European (B) and the North Sea (C) lineages (Fig. 3; Feng et al. 2020).

### Patterns of genetic parallelism

When including relatedness as a random effect in association tests between SMLAs and ecotype (EMMAX), *λ* was reduced from *λ*=1.95 to *λ*=1.01 and from *λ*=1.73 to *λ*=1.32 for three- and nine-spined sticklebacks, respectively. Thus, accounting for relatedness reduced *p*-value inflation completely in three-spined sticklebacks, but not in nine-spined sticklebacks. Nevertheless, GC ensured that any residual *p*-value inflation was accounted for except when using permutation. After corrections for *p*-value inflation and multiplicity, the number of outlier regions that were significant in at least 5% of all parameter combinations/correction methods (*C*≥0.05) was 26 for three-spined sticklebacks and nine for nine-spined sticklebacks (Fig. 4 & 5). While no single parameter combination detected all these outlier regions the most successful LDna parameter settings for three-spined sticklebacks were *|E|_min_*=10, *Cor_th_*=0.5 and any combination of *SNP_min_* =[10, 20] (detecting all but the two ChrIX outlier regions) with the corresponding parameter settings for nine-spined sticklebacks being *|E|_min_* =20, *SNP_min_*=10 and any combination of *Corth=*[0.6, 0.7, 0.8] (detecting all but the ChrXIV outlier region). However, no outlier regions in nine-spined sticklebacks were found in >50% (*C*>0.5) of the parameter combinations, while this was the case for seven outlier regions in three-spined sticklebacks (Fig. 5; Supplementary Table 1), with the corresponding numbers for *C*>0.25 being 13 and four, respectively. Thus, regardless of *C*-score, the number of outlier regions detected were always larger in three-than in nine-spined sticklebacks, showing that outlier detection in nine-spined sticklebacks was more dependent on parameter settings than that in three-spined sticklebacks. As a consequence, widely different results could have been obtained, particularly for nine-spined sticklebacks, if only a single (arbitrary) parameter setting would have been chosen for LDna.

**Figure 4.**
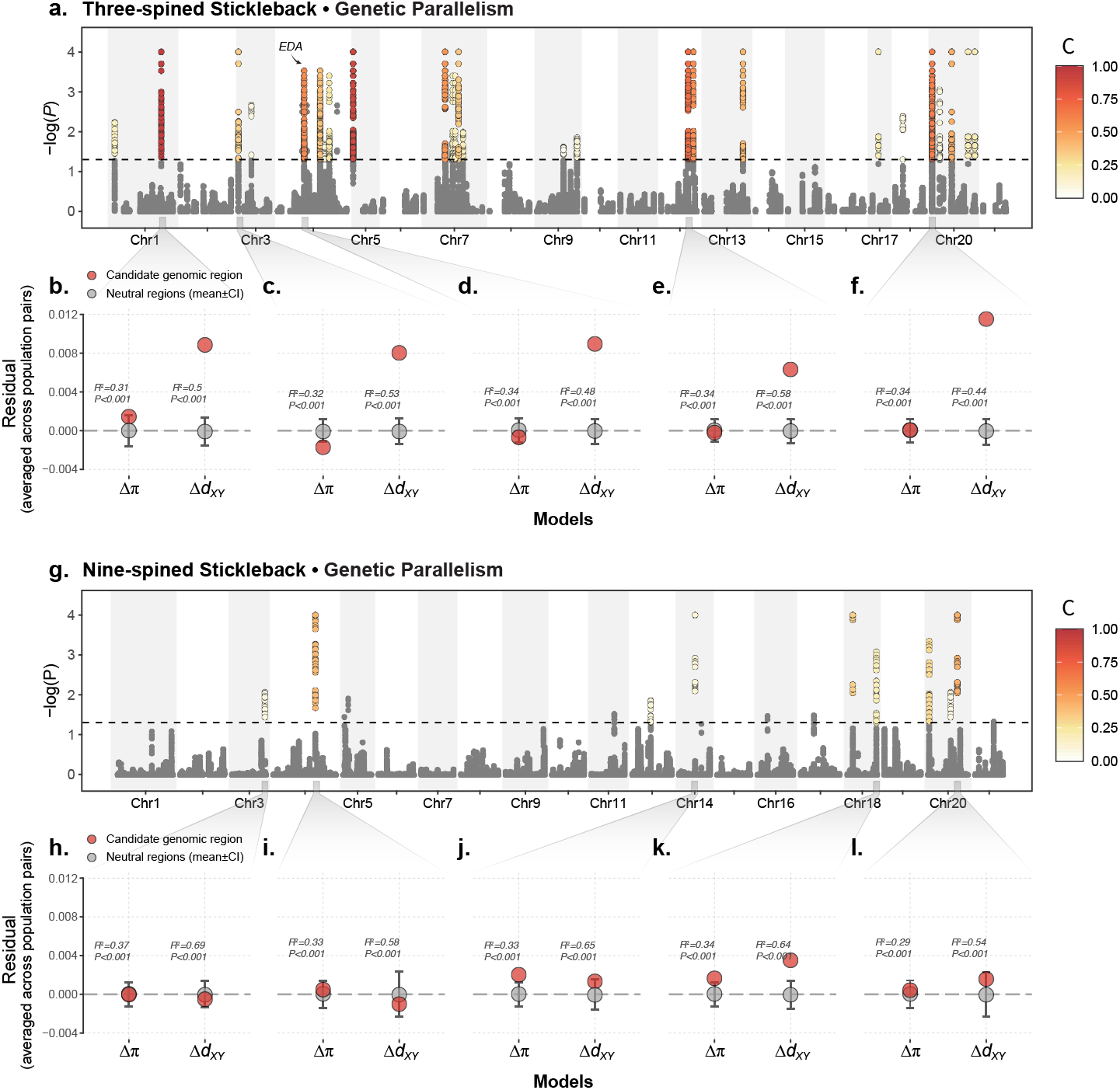
Three-spined sticklebacks exhibit stronger level of parallel genetic evolution than nine-spined sticklebacks. (a,g) Manhattan plots of −log10(*P*) value testing for associations between LD-clusters and ecotype (EMMAX). Colour for each unique genomic region indicates the proportion of all combinations parameter/correction method settings a given region was found significant in (*C*-score) after corrections for multiple testing and *p*-value inflation (grey indicates *C*-score<0.05). (b-f, h-l) summaries of residuals of linear regression models based on the genetic diversity *(π*) and genetic divergence *(d_XY_*) derived from marine-freshwater population pairs for selected outlier regions (see Materials and methods). The squared correlation coefficient (*r*^2^) is shown as an averaged value across all models from different population pairs. Summaries for all regions are given in the Supplementary Fig. 5. All models were statistically significant (*P*<0.001).

While all outlier regions with high *C*-scores tended to also have large effect sizes (Fig. 5), some outlier regions with large effect sizes (e.g. Chr17_1, Chr20_4 and Chr20_5 for three-spined sticklebacks and Chr14_1 for nine-spined sticklebacks) did not have high *C*-scores (Fig. 5). These regions were more sensitive to parameter settings, but when they were detected, they tended nevertheless to have large effect sizes (and be highly significant). The Chr14_1 (*C*=0.11) outlier region in nine-spined sticklebacks, for instance (with the cor=0.45 [95% quantile for all LD-clusters mapping to the region] both when correcting and not correcting for relatedness), was only detected when *|E|_min_*=40 and *SNP_min_*=40.

**Figure 5.**
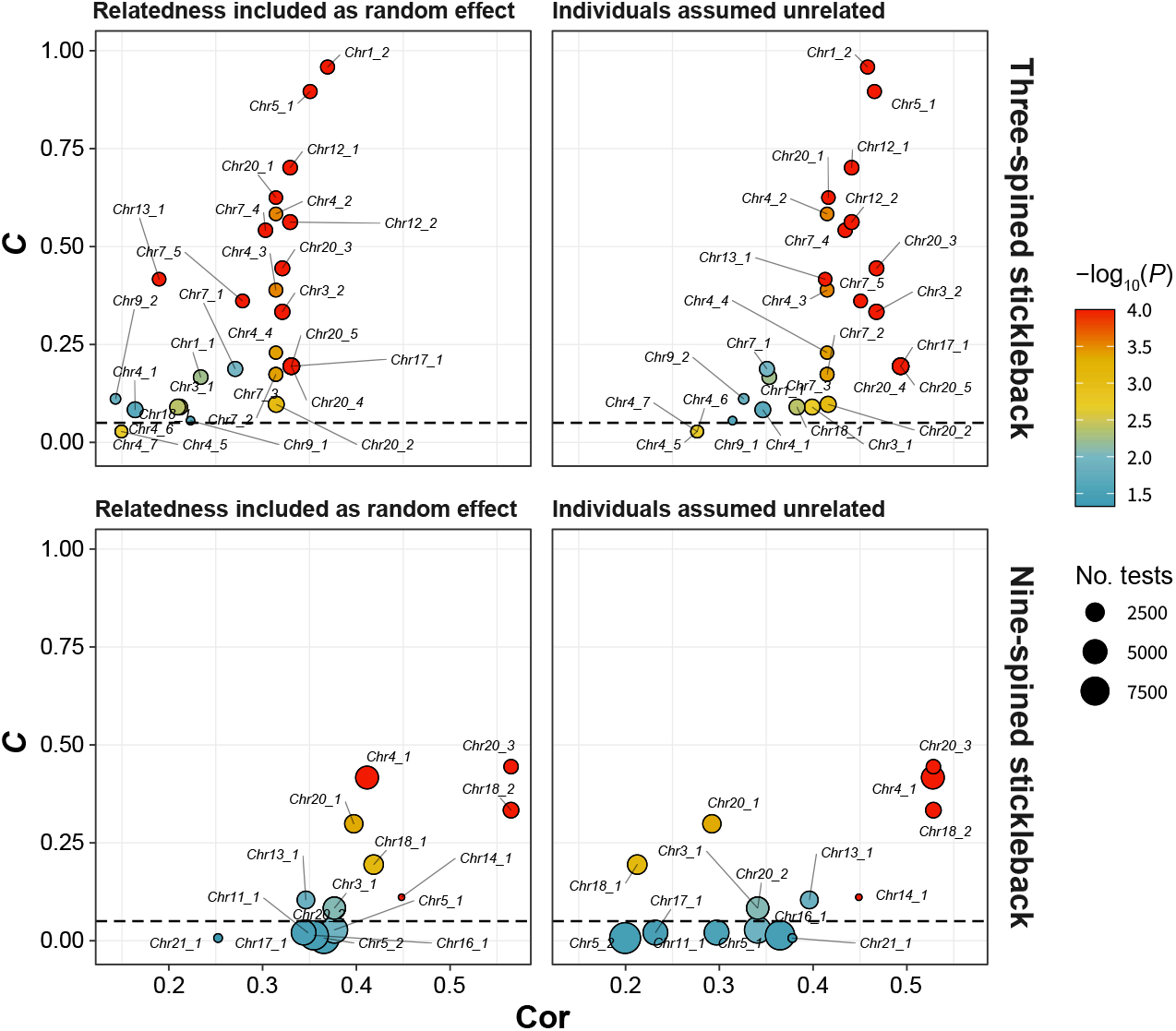
Relationship between C-score and effect size. Figure depicts *C* as a function of effect size (*cor*; 95% quantile across all significant LD-clusters mapping to a given genomic region) for outlier genomic regions, when (left panel) relatedness is accounted for and (right panel) when individuals are assumed to be unrelated for three- and nine-spined sticklebacks (upper and lower panels, respectively). Size indicates the mean number of tests for significant LD-clusters for a given region (a function of *|E|_min_, SNP_min_* and *Cor_th_*, see main text for details), with smaller numbers indicating that a region is only significant when fewer and larger LD-clusters are tested and colour indicates the most significant P-value across any correction method (“A+fdr”,”A+perm”, “GC+fdr” and “perm”). Only outlier regions above the horizontal dashed line (*C*>0.05) are considered in our analyses. 3sp, three-spined stickleback; 9sp, nine-spined stickleback.

Focusing on four different geographic regions within the Atlantic Ocean, the mean effect size (estimated based SMLAs from all significant loci from a given outlier region) across all genomic regions was *cor_unrl_*=0.38 (sd=0.24) and *cor_A_*=0.28 (sd=0.17) for three-spined sticklebacks and *cor_unrl_*=0.29 (sd=0.20) and *cor_A_*=0.27 (sd=0.17) for nine-spined stickleback. In both species, no outlier region associated with marine-freshwater parallelism displayed universally high effect sizes across all analysed geographic regions (Fig. 6). The geographic region with the highest mean effect size across the outlier regions for three-spined sticklebacks was the Norwegian Sea *(cor_unrl_*=0.38, sd=0.17; *cor_A_*=0.66, sd=0.2), with least evidence for parallelism being found in White & Barents Sea *(cor_unrl_*=0.24, sd=0.15; *cor_A_*=0.14, sd=0.078) and in the Baltic Sea regions *(cor_unrl_*=0.294, sd=0.19; *cor_A_*=0.22, sd=0.14). In contrast, the two geographic regions with highest effect sizes across all outlier genomic regions in nine-spined sticklebacks were the opposite of three-spined sticklebacks, namely White & Barents Sea *(cor_unrl_*=0.43, sd=0.17; *cor_A_*=0.39, sd=0.17) and the Baltic Sea regions *(cor_unrl_*=0.41, sd=0.22; *cor_A_*=0.42, sd=0.19).

**Figure 6.**
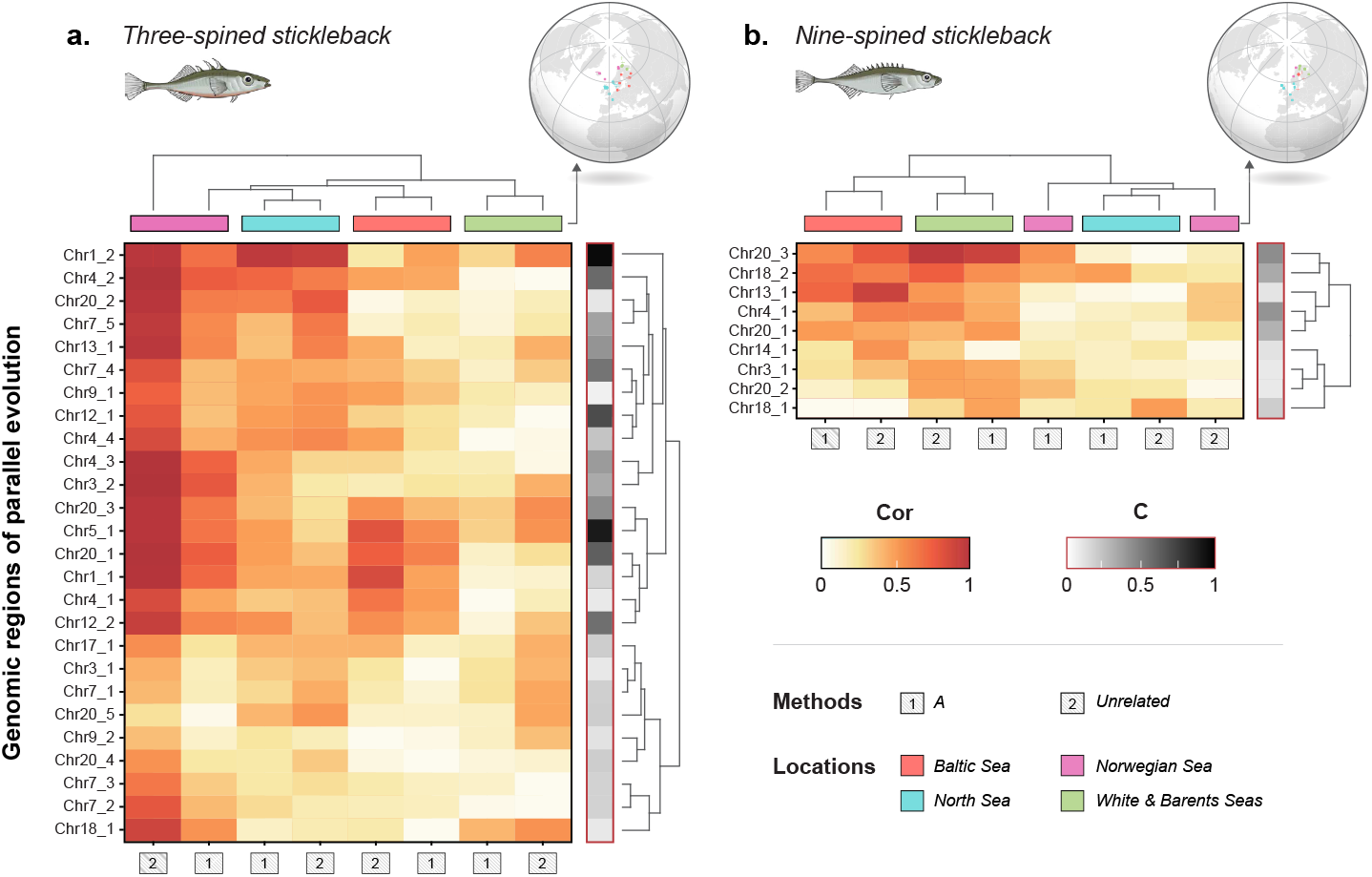
Regional parallelism. Shown are heatmaps of effect sizes (*cor*) from linear regressions between SMLAs and ecotype (EMMAX) for outlier regions (*C*-score≥0.05) separately for four geographic regions. Results are shown for (a) three- and (b) nine-spined sticklebacks, when assuming individuals are unrelated (unrl) and when a relatedness matrix (A) was included as a random effect. Grey side bars indicates the *C*-score.

The mean size for outlier regions was 3.0 times larger for three-spined sticklebacks compared to nine-spined sticklebacks (154Kbps *vs.* 52Kpbs; Supplementary Table 1), with the mean number of unique significant loci mapping to each region being 2.7 times larger in three-compared to nine-spined sticklebacks (115 SNPs with a range of 10-1341 *vs.* 42 SNPs with a range of 10-109). The total number of loci from significant LD-clusters mapping to outlier regions was 379 and 2996, for three-spined and nine-spined sticklebacks, respectively. However, a large proportion of the loci (43%) in three-spined sticklebacks mapped to the Chr1 inversion.

The genomic regions under parallel evolution in three-spined sticklebacks show an excess in absolute divergence (Δ*d_XY_*) compared to neutral genomic regions, suggesting ancient origin of the selected regions (as per Nelson & Cresko 2018). On the contrary, the genomic regions under parallel evolution in nine-spined sticklebacks do not show significant *Δd_XY_*. Specifically, in three-spined sticklebacks, 16 out of 26 genomic regions under selection had higher *Δd_XY_* than the 95% CI of *Δd_XY_* estimated from 100 random “neutral” regions across the genome l (Fig. 4a-d, Supplementary Fig. 8). Those ancient regions included EDA gene and the inversion in Chr1 (Jones et al. 2012; Fang et al. 2020a; Fig. 4d,b). In contrast, only one out of nine regions under selection in nine-spined sticklebacks showed (slightly) higher Δ*d_XY_*, approaching the 95% CI of Δ*d_XY_* for neutral regions (Fig. 4k). Full results of genetic diversity and divergence in the candidate genomic regions were given in Supplementary Fig. 8.

There was a significant negative correlation between divergence times and the level the genetic parallelism in nine-spined sticklebacks (MLPE: *β*=−2e-4, P<0.001; Supplementary Fig. 9). However, there was no significant correlation in three-spined sticklebacks (MLPE: *β*=−0.014, *P*=0.76; Supplementary Fig. 9).

## Discussion

Theory predicts the probability of parallel evolution to be negatively correlated with divergence time and positively correlated with effective population size and population connectivity (MacPherson & Nuismer 2017). Despite their similar life histories, ecologies and distribution ranges, three- and nine-spined sticklebacks show dramatically different phylogenetic histories, within population genetic diversities and population structuring across comparable geographic distances and as predicted, very different levels of genetic parallelism in response to colonisation of freshwater environments. As the results show, gene flow between nine-spined stickleback populations is more restricted than between three-spined stickleback populations, resulting in a more heterogeneous pool of SGV available for freshwater adaptation, thereby reducing the probability of parallel evolution in the former. These findings indicate that the two species differ markedly in the fundamental processes affecting the distribution of adaptive genetic variation among their demes.

Three-spined sticklebacks displayed a larger number (2.9 times) of genomic regions involved in marine-freshwater parallelism than nine-spined sticklebacks, and these regions were on average three times larger and harboured eight times more loci. Most outlier regions in three-spined sticklebacks show excess divergence (△*d_XY_*), a result in line with evidence from other studies (Nelson & Cresko 2018) suggesting ancient origins of haplotypes involved in parallel evolution in this species. In contrast, the outlier regions in nine-spined sticklebacks do not follow the same pattern (i.e. no significant △*d_XY_*). Furthermore, for both species, no marine-freshwater divergence associated region showed high effect sizes across all studied geographic regions, suggesting that parallelism is often geographically limited. Below we discuss how these findings shed new light into our understanding of how geographic patterns of parallel local adaptation is shaped by demographic and phylogenetic history.

### Genetic diversity, population connectivity and phylogenetic history

Genetic diversity was generally lower for nine-than three-spined sticklebacks and in freshwater compared to marine populations. These findings align with earlier studies showing lower genetic diversity in nine-than in three-spined stickleback populations (DeFaveri et al. 2012; Merilä 2013), and in freshwater fish populations than marine fish populations in general (Avise et al. 1987; DeWoody & Avise 2000; McCusker & Bentzen 2010; Ward et al. 1994; Ward et al. 1992). Assuming that three-and nine-spined sticklebacks exhibit similar mutation rates, the observed differences in *θ* (three-spined > nine-spined stickleback) should reflect differences in coalescent *N_e_* (as *θ =4N_e_μ).* Hence, lower *N_e_*, stronger population structure and stronger IBD all contribute to more heterogeneous pools of SGV in nine-spined than in three-spined sticklebacks. Assuming that adaptive genetic variation follows the same general pattern, SGV for freshwater adaptation is expected to be considerably reduced in nine-compared to three-spined sticklebacks.

Our analyses revealed that the nine-spined stickleback lineages were far older than three-spined stickleback lineages, and that the degree genetic parallelism decreased as an increasing function of divergence time in the nine-spined stickleback. All this supports the notion that differences in divergence time influence the probability of parallel evolution, as pools of SVG get increasingly differentiated with time (MacPherson & Nuismer 2017). However, the question of which is a more important determinant of the probability of parallel evolution – divergence time or gene flow – is not easily answered because the two are not independent: restricted gene flow is a prerequisite for the formation and maintenance of distinct lineages. It is therefore difficult to disentangle the effect of current population structure and past divergence on the levels of genetic parallelism because both are functions of the species demographic history.

### Geogra/hic heterogeneity in selection optima?

Parallel evolution not only requires access to the same pool of SGV (Barrett & Schluter 2008; Schluter & Conte 2009), but also parallelism of selection optima across the distribution range, which cannot be taken for granted (Bolnick et al. 2018; Magalhaes et al. 2020; Stuart et al. 2017). Thus, lower parallelism in selection optima across freshwater habitats could also explain the lower degree of parallelism in nine-than three-spined sticklebacks. However, geographic differences in SGV can result in contrasting patterns of genetic parallelism, both globally (Fang et al. 2020a) and locally (Leinonen et al. 2012). Based on a small subset of the data used here, simulations in Kemppainen et al. 2021 show that the level of IBD characteristic of nine-spined stickleback populations (as opposed to three-spined stickleback-like scenarios) is sufficient to severely restrict SGV for local adaptation. Thus, while parallel angles of selection are a prerequisite, parallel evolution also relies on access to the necessary ancestral SGV for local adaptation. That ancestral SGV in turn is determined by population demographic parameters such as *N_e_* and population connectivity.

The reported differences in the degree of parallel evolution between two stickleback species could be argued to be an artefact of the inherent difficulty of detecting outlier loci among highly differentiated populations (Galloway et al. 2020; Hoban et al. 2016; Matthey-Doret & Whitlock 2019). However, whenever background differentiation is high (due to population structuring), we can also expect *p*-value inflation due to relatedness. Accounting for relatedness can in some circumstances be expected to increase statistical power to detect outliers (Kang et al. 2010; Kang et al. 2008). For instance, when multiple populations in similar habitats display high frequencies of the same genetic variants (i.e. parallel evolution); the more divergent the populations are, the stronger the contrast between neutral genetic background (reflecting relatedness) and genomic regions under selection will be. Since neither effect sizes nor *p*-value inflation (both of which are important determinants of statistical power) differed much between nine-and three-spined sticklebacks for the outlier regions, there is no reason to doubt the conclusion that the marine-freshwater differentiated regions in three-spined sticklebacks outnumber such regions in nine-spined sticklebacks across a wide range of parameter/threshold values and correction methods. This and other methodological considerations are discussed further in Supplementary Information 2.

### Geogra/hic heterogeneity in marine-freshwater parallelism

Historically, much of the parallel evolution research in three-spined sticklebacks has focused on Eastern Pacific populations (reviewed in Fang et al. 2020a). However, it is becoming increasingly clear that parallelism in three-spined sticklebacks is not as universal and global as previously thought (Fang et al. 2020a). This is also clear from our analyses (with more extensive geographic coverage than in previous studies), where heterogeneity in effect sizes across the different geographic region was found not only in nine- (where this was expected) but also in three-spined sticklebacks. In our analyses of genetic parallelism, we limited our comparisons to the Atlantic region as we lacked samples of marine nine-spined sticklebacks from the Eastern and Western Pacific regions. Therefore, we do not know if the patterns seen in the Atlantic region can be generalized to the rest of the species distribution range. However, there is a good reason to believe that inclusion of populations from the Eastern Pacific would have revealed stronger, not weaker, differences in levels of parallel evolution between the two taxa. Namely, the extent of genetic parallelism in the three-spined stickleback from the Pacific region is far stronger than that in the Atlantic (Fang et al. 2020a). While this might at first suggest that inclusion of Eastern Pacific samples to the comparison might recover more genetic parallelism also in the nine-spined stickleback, we believe the opposite is more likely to be true. The Eastern Pacific is the ancestral range of the three-spined stickleback, which harbours most of the SGV involved in marine-freshwater adaptation (Fang et al. 2020a; Fang et al. 2020b; Fang et al. 2018). In contrast to three-spined sticklebacks, the oldest populations of nine-spined sticklebacks are located in the Western Atlantic region (Fig. 3), and therefore, it is logical to assume that at least a part of the SGV involved in parallel evolution might have been lost during the invasion of the Pacific region. Hence, the inclusion of Pacific populations of both species into the analyses would likely reveal even stronger differences than observed now.

### Age of freshwater adapted alleles

While nine-spined stickleback populations have a longer evolutionary history than the three-spined stickleback populations, the genomic regions under parallel selection in three-spined sticklebacks appear to be of more ancient origin than the populations in which they are segregating. A possible explanation for this counterintuitive finding may lie in the effect of gene flow and *N_e_* on the ability of species maintain ancestral haplotypes in the pool of SGV; the larger *N_e_* and the weaker population subdivision makes this scenario more likely in three-compared to nine-spined sticklebacks. Although the Atlantic three-spined stickleback populations have a relatively young evolutionary history (colonisation occurred 29.5-226.6 Kya; Fang et al. 2020b), the freshwater-adapted alleles in the SGV pool were inherited originate from Eastern Pacific region (Fang et al. 2020a), which harbours these ancient t haplotypes (~six million years old; Nelson & Cresko 2018). In contrast, lower *N_e_* and gene flow in the nine-spined stickleback marine populations is expected to limit the maintenance and geographic spread of SGV. This could lead to higher turnover rates of freshwater adapted alleles, higher dependence on novel mutations and consequently higher geographic heterogeneity in parallel evolution in nine-spined sticklebacks.

### Implications for local adaptation

The implications of the absence of a homogenous pool of SGV in the nine-spined stickleback are relevant to local adaptation in general. Populations colonizing new environments may lose potentially beneficial variation via bottlenecks and founder events. With limited or absent gene flow, the lost variability cannot easily be regained. Consistent with this, Kemppainen et al. 2021 demonstrated that the genetic architectures underlying pelvic reduction (a common freshwater adaptation in sticklebacks) is highly heterogeneous in nine-spined sticklebacks even across short (<10 km) geographic distances. In addition, many freshwater populations lacked pelvic reduction altogether probably because they lacked the SGV underlying this trait, thus restricting the populations to suboptimal phenotypes. A possible example where restricted access to SGV has led to potentially less optimal freshwater adaptation can also be found in three-spined sticklebacks. In a small and isolated region in the northern Finland, fully plated three-spined sticklebacks displayed reduced lateral plate height possibly as a compensatory adaptation to the genetic constraint imposed by the lack of the low-plate EDA allele in these populations (Leinonen et al. 2012). Thus, it is important to note that lack of large pool of SGV not only limits parallel evolution, but also local adaptation more generally.

The lack of SGV to fuel local adaptation can be mitigated by introgression between divergent clades as this can substantially increase the genetic variation in the affected populations (Anderson 1949; Arnold 1997; Marques et al. 2019). This appears to be the case for the Baltic Sea nine-spined stickleback populations which have experienced introgression from divergent western European populations (Feng et al. 2020; Shikano et al. 2010b; Teacher et al. 2011); a comparison of admixed and non-admixed populations revealed the former to have significantly higher heterozygosity than the latter. Since admixed populations were excluded from the analyses of within population genetic diversity, this did not have any effect on our inference beyond reducing sample sizes available for analyses. Nevertheless, the results demonstrate that introgression is an important determinant of genetic diversity, and thus potentially also of the adaptive potential of populations. This is consistent with Baltic Sea having large effect sizes for a subset of the regions that have large effect sizes also in the White & Barents Sea possibly compensating for the lack of freshwater adaptations in the Western lineage (which have much lower effect sizes for all outlier regions).

### Genetic differentiation and speciation

It is intriguing that the two species of sticklebacks studied here do not display only highly contrasting population structures but come from different genera containing an equally contrasting number of recognized species. While it is tempting to think that there could be a causal connection between among population connectivity and propensity to speciate, one needs to notice that there is also evidence to suggest that there may be yet undescribed species in both genera (Guo et al. 2019; Taylor et al. 2006). Hence, counting taxonomically valid species as proxies of speciosity can be misleading.

What may be more interesting to consider is the process of speciation. In the case of the three-spined stickleback “species pairs”, speciation seems to progress rapidly towards completion, but full reproductive isolation is not usually reached (McKinnon & Rundle 2002). In fact, whenever the ecological conditions that drove the evolution of reproductive isolation in the first place cease to exist, hybridization and reverse speciation is known to occur (Marques et al. 2019; Rudman & Schluter 2016). Hence, this suggests that strong genetic incompatibilities have not had time to evolve. The marked between species difference in divergence times of local three- and nine-spined populations shown in this study suggest that one should expect speciation through accumulation of genetic incompatibilities to be much more likely in the *Pungitius* than in the *Gasterosteus* genus. There is indeed some evidence for evolution of incompatibilities in *Pungitius* (Natri et al. 2019), and although hybridization has occurred fairly frequently (Guo et al. 2019), the species have not collapsed into hybrid swarms. It is tempting to speculate that the same features that can drive local adaptation and initial progress along the speciation continuum in the short term are actually limiting factors in driving speciation to completion. Hence, the two genera should provide an interesting model system for future studies focused on the roles of adaptive divergence *vs.* incompatibilities in generating new species.

In conclusion, the results establish that the two co-distributed stickleback species possess strikingly different population genetic structures suggesting far more limited gene flow, and hence, a more heterogeneous pool of SGV among the nine-spined, than among the three-spined stickleback populations. This greater heterogeneity likely underlies the observed lower degree of genetic parallelism among the nine-spined stickleback populations. Furthermore, there appears to be generally less SGV in the nine-than in the three-spined stickleback, possibly because of lower long-term effective population sizes of the latter. However, high levels of SGV were detected in those few nine-spined stickleback populations where introgression from other related lineages or species has been documented. All in all, the results suggest that because of the contrasting levels of heterogeneity in SVG, the two stickleback species are differently disposed to adapt similar selection pressures via parallel and non-parallel genetic mechanisms.

## Acknowledgements

We thank members of EGRU for very useful feedback and discussions during the manuscript preparation, and Sami Karja, Kirsi Kähkönen, Laura Hänninen and Miinastiina Issakainen for help with the laboratory work and its planning. We are also grateful for all the people who helped in sampling and obtaining the nine-spined stickleback samples used in this study, including: Michael Bell, Victootr Berger, Pär Byström, Jacquelin De Faveri, Lasse Fast Jensen, Abigel Gonda, Gábor Herczeg, Kjetill Hindar, Nellie Konijnendijk, Dmitry Lajus, Tuomas Leinonen, Vladimir Loginov, Scott McCairns, Andrew McColl, Henri Persat, Tom Pike and Joost Raeymaekers. Our research was support by Academy of Finland (grants # 129662, 134728 and 218343 for JM; 1316294 to PM), Helsinki Institute of Life Sciences (HiLife for JM), Finnish Cultural Foundation (00190489 to PK), and Chinese Scholarship Council grant (201606270188) to BF.

