## Supplementary materials for "Population structure limits parallel evolution"

### Content

#### Supplementary Methods

**Supplementary Method 1 |** Genotype likelihood estimation.

**Supplementary Method 2 |** Genetic diversity and differentiation.

**Supplementary Method 3 |** Comparative phylogenomic analyses.

**Supplementary Method 4 |** Complexity reduction using nested three-step linkage disequilibrium network analyses (LDna).

**Supplementary Method 5 |** Testing associations between “synthetic alleles” from LD-clusters and ecotypes using linear mixed models.

**Supplementary Method 6 |** Restricted maximum likelihood (REML) analyses.

**Supplementary Method 7 |** Correction for multiple testing by permutation.

**Supplementary Method 8 |** Genetic diversity and divergence in the marine–freshwater divergent genomic regions.

#### Supplementary Figures

**Supplementary Figure 1 |** Sampling maps of the populations used in different analyses.

**Supplementary Figure 2 |** Nested three-step LDna complexity reduction.

**Supplementary Figure 3 |** The relationship between EMMAX analyses performed on “synthetic alleles” from PCA-analyses of LD-clusters.

**Supplementary Figure 4 |** Cumulative numbers of outlier regions detected for randomly selected parameter/correction method combinations.

**Supplementary Figure 5 |** The maximum-clade-credibility summary trees of the SNAPP phylogenies reported in Fig. 3, main document.

**Supplementary Figure 6 |** Supplementary genetic diversity estimates.

**Supplementary Figure 7 |** Intra-clade genetic differentiation.

**Supplementary Figure 8 |** Genetic diversity and divergence in the genomic regions under selection.

**Supplementary Figure 9 |** Correlations between divergence time and genetic parallelism.

#### *Supplementary Tables*

**Supplementary Table 1 |** Detailed results from regional EMMAX analyses.

**Supplementary Table 2 |** Information on the nine-spined stickleback samples used in the study.

**Supplementary Table 3 |** Populations in Figure 1b, c.

**Supplementary Table 4 |** Information on LD-clusters of three- and nine-spined sticklebacks.

#### *Supplementary Information*

**Supplementary Information 1 |** Outlier population (RUS-LEV) in the IBD analyses.

**Supplementary Information 2 |** Methodological considerations.

#### 1 | Genotype likelihood estimation

In the step of quality filtering in ANGSD, we retained only bases with a minimum Phred score of 20 and a minimum mapping quality of 25 (-minQ 20 and -minMapQ 25). We retained sites with less than 20% missing data and a minimum read depth of two for each individual following Fang et al. 2020 to account for low sequence-depth data and computed genotype likelihoods for variants that had a *p*-value smaller than 1e-6 (-SNP_pval 1e-6). The sex chromosomes were excluded (chromosome 19 [group XIX] in the three-spined stickleback; Kitano et al. 2009; Natri et al. 2013; chromosome 12 [LG 12] in the nine-spined stickleback; Natri et al. 2019; Rastas et al. 2015; Shapiro et al. 2009). The full scripts of ANGSD pipelines can be found from Fang et al. 2020 (https://doi.org/10.5061/dryad.b2rbnzsb1).

#### 2 | Genetic diversity and differentiation

For *p* and *θ*, site allele frequency likelihoods and the site frequency spectrum (SFS) were estimated for each population in ANGSD. The summary statistics of *p* and *θ* for each population were calculated using a custom R scripts from Momigliano et al. 2021. Scripts can be found in: https://github.com/Nopaoli/Demographic-Modelling/blob/master/Diversity_fromSFS/Stats_from_SFS_TD.R. To calculate *H*, we first estimated the SFS for each individual and divided the number of segregating sites (the second entry of the SFS) by the sum of the SFS. Population estimates of *p* and *θ* were only applied to the populations consisting of multiple individuals (sample size n ≥ 2). The samples from Jones et al. 2012 in the dataset of three-spined sticklebacks (c.f. Fang et al. 2020) were excluded in the *H* calculation due to low read depth (≤ 1). The Supplementary Fig. 1 presents the sampling maps of the populations used in different analyses.

Since some sampled populations are known to be admixed (Feng et al. 2020; Guo et al. 2019), their genetic diversity was expected to be elevated. Therefore, the analyses of *H*, *p* and *θ* were conducted both excluding and including the known admixed nine-spined stickleback populations (information from Guo et al. 2019 and Feng et al. 2020), and both results were reported (Fig. 1 and Supplementary Fig. 2). Specifically, seven out of 29 freshwater populations (BEL-MAL, DEN-RES, PP-FR-MAR, PP-RU-BLS, PP-RU-GOR, PP-RU-KVA, PP-RU-MAG) were identified to be admixed with other *Pungitius* species (Guo et al. 2019), and six out of seven marine populations (GER-RUE, FIN-HEL, SWE-BOL, FIN-KIV, DEN-NOR, SWE-FIS) have experienced varying degree of introgression between divergent *P. pungitius* lineages (Feng et al. 2020).

To calculate $F_{ST}$ in each species, genotypes were called in ANGSD (-doVcf 1) with additional filtering on the genotype posterior probability (-postCutoff 0.95). VCFtools v.0.1.15 (Danecek et al. 2011) was used for the downstream SNP filtering, setting maximum missing genotypes to 20% (--max-missing 0.8), a minimum minor allele frequency to 0.05 (--maf 0.05) and minimum distance between any two sites to 30,000 bp (--thin 30,000). We then used this subset of high quality, called genotypes to estimate global $F_{ST}$ for each ecotype and the pairwise $F_{ST}$ between all populations with the R packages *hierfstat* (Goudet 2005) and *StAMPP* (Pembleton et al. 2013), respectively.

#### 3 | Comparative phylogenomic analyses

For both species we filtered SNPs called from ANGSD in VCFtools, retaining only "unlinked" (i.e. no closer than 10 kb; --thin 10000) bi-allelic loci (--min-alleles 2; --max-alleles 2) with no missing data (--max-missing 1) and a minor allele frequency above 0.01.

For nine-spined sticklebacks, the first calibration was the time to the most recent common ancestor (TMRCA) of all lineages, 2.69 Mya (million years ago, 95% Highest Posterior Density [HPD] interval: 1.87–3.57 Mya). The second was the TMRCA of European lineages at 0.67 Mya (95% HPD interval: 0.43–0.92 Mya). For three-spined sticklebacks, the first calibration was the TMRCA of all lineages, 191.7 Kya (thousands of years ago; mean of the time estimates in Fang et al. 2020b [36.9–346.5 Kya]). The second was the divergence time between the Pacific and Atlantic lineages, 128.1 Kya (29.5–226.6 Kya). The third one was the TMRCA of the Northern European lineages, 53.25 Kya (11.3–95.2 Kya). The calibration points are visualized in the Supplementary Fig. 3.

To perform SNAPP analyses, the input XML file was prepared using a Ruby script "snapp_prep.rb" (github.com/mmatschiner/snapp_prep). Stationarity and convergence of 100 million Markov-Chains Monte Carlo (MCMC) from SNAPP were checked in TRACER v1.6 (ESS > 1000; beast.bio.ed.ac.uk/tracer/). A maximum-clade-credibility summary tree was generated using TreeAnnotator (Drummond et al. 2012). The SNAPP trees were visualised by the program DensiTree v.2.2.6 (Bouckaert 2010) with a 10% burn-in for each MCMC chain.

#### 4 | Complexity reduction using nested three-step linkage disequilibrium network analyses (LDna)

Since allelic states of many pairs of loci can be highly correlated (in linkage disequilibrium, LD), corrections for multiple testing – assuming all loci are independent – are unnecessarily conservative. Recently, Li et al. 2018 (see also: Santure & Garant 2018) developed a method where correlated sets of loci (LD-clusters) along non-overlapping windows along chromosomes can be grouped together by LD network analyses (LDna; Kemppainen et al. 2015) and summarised by Principal Component Analyses (PCA) prior to Genome Wide Association (GWA)-analyses. Furthermore, in the presence population structuring, LD typically extends across the whole genome such that complexity reduction can be increased further by performing LDna within chromosomes or even at the whole genome level (prior to GWA-analyses). The idea is that for all (genome wide) loci that only reflect population structuring (thus cannot be reliably associated with any of the studied traits due to relatedness) only one test is sufficient, thus substantially reducing the total number of tests needed. The benefit of this, relative to window-based approaches that disregards the correlation structure among loci, is the possibility to separate different sets of correlated loci even though they are interspersed in the genome (Li et al. 2018), as demonstrated in Supplementary Figure 2a.

Since LD-estimates between all pairs of loci from large population genomic data set (with millions of loci) cannot be considered simultaneously, we adopt a nested three-step approach to perform complexity reduction (Fang et al. 2020; Li et al. 2018) prior to downstream analyses. In LDna step 1 (LDna-1) we recursively identified LD-clusters along chromosomes where at least one locus was connected by LD>0.7 with all other loci in its cluster. These SNPs represent the whole cluster in the subsequent analysis steps (rSNPs; if several were found, one SNPs was randomly chosen; Supplementary Fig. 2a). To reduce computational speed this recursive algorithm was performed within non-overlapping windows of 1000 bi-allelic SNPs and within those windows, LD was estimated in windows of 100 SNPs with minor allele frequency, *maf*>0.05, (Fang et al. 2020). Thus, while no SNP could directly be connected to another SNP further than 100 SNPs away, they could nevertheless belong to the same cluster via links to other (intermediate) SNPs, and SNPs from different (non-overlapping) 1000 bps windows can of course become connected in LDna-2. The median proportion of the genetic variation explained by PC1 for such LD-clusters typically exceeds 99.9%.

In the second step (LDna-2; Supplementary Fig. 2b), using the rSNPs from LDna-1, we performed LDna separately for each chromosome ignoring all singleton clusters (SNPs that were not connected to any other locus by at least LD = 0.7), as in Fang et al. 2020. In standard LDna (Kemppainen et al. 2015), all pair-wise LD values are considered simultaneously using single-linkage clustering trees, where each cluster is defined by its weakest LD-value among any pair of loci in the cluster. LD-clusters of interest are defined by two parameters; the minimum number of edges in the cluster ($|E|_{min}$, which is sensitive to both the connectedness within the cluster as well as the number of loci in a cluster) and a threshold for λ ($\lambda_{lim}$), which indicates the magnitude of how different LD-clusters are with respect to median LD when considered separately or jointly (Kemppainen et al. 2015). However, similar cluster solutions that can be obtained by different combinations of $|E|_{min}$

and $\lambda_{lim}$ can also be obtained just by varying $|E|_{min}$. Since these parameters are redundant, we here considered all branches for a given $|E|_{min}$ as separate LD-clusters regardless of λ (thus removing this parameter altogether) as shown in Supplementary Fig. 2b. Since, for a given value of $|E|_{min}$, PC1 in “LD-clades” with node depths ≥0.8 (i.e. when the lowest LD value between any two loci in the cluster is ≥0.8) typically explain >>90% of the variation, these were considered as single LD-clusters; such cases increase with lower $|E|_{min}$ as this typically leads to more branching at higher LD levels. In previous implementations of the three-step LDna (Fang et al. 2020), the most interconnected SNP (the SNP with the highest median LD with all other loci in its cluster) in each LD-cluster/branch from LDna-2 was chosen as the rSNP for LDna-3. However, since the genetic variation in a cluster with many loci more efficiently and robustly can be summarised by PC-coordinates (based on all the loci in the cluster), we here opted to base the LDna-3 tree on all pairwise correlation coefficients of PC1 coordinates between the LDna-2 LD-clusters entering LDna-3 (Supplementary Fig. 2c). The matrix of all pair-wise correlation coefficients $K_{PC1}$ were nevertheless always correlated with the corresponding matrix of LD $K_{LD}$ based on rSNPs, so this is not likely to substantially affect the final LDna-analysis, except to make it less sensitive to the exact choice of the rSNP.

*Data sets*

To make a rigorous comparison, the comparative LDna analyses between three- and nine-spined sticklebacks were restricted to the Atlantic region. This is because our sampling lacked marine nine-spined stickleback populations from the Pacific regions and because three-spined sticklebacks from the Eastern Pacific region exhibit unusually high (ca. 10 times) genetic parallelism compared to other regions in the world (Fang et al. 2020).

To generate comparable datasets for three- and nine-spined sticklebacks for LDna and downstream analyses, it is important to consider that genome coverage was much higher in nine- than three-spined sticklebacks due to larger proportion of WGS samples in the former species (80.1%) than in the latter species (22.9%). Therefore, firstly, we made a list of sites across the genome for both species for which we had sufficient coverage of high-quality reads using ANGSD (-minIndDepth 1, -uniqueOnly 1, -remove_bads 1, -minMapQ 20, -minQ 20, -minInd 25%), resulting in 101,598,319 and 129,356,224 sites in three- and nine-spined stickleback genomes, respectively. Then we retained variable sites using the -SNP_pval flag in ANGSD (-SNP_pval $1e^{-6}$) and filtered by minor allele frequency (maf ≥ 0.05) using custom R scripts based on the output ‘GENO’ file. After this, 882,125 SNPs and 1,725,617 SNPs were retained from three- and nine-spined sticklebacks, respectively. Next we computed the number of SNPs that we would have discovered in nine-spined sticklebacks if we had sequenced the same number of bases as for the three-spined sticklebacks: $SNPs_{9s}=SNPs_{3s}/L_{3s}*L_{9s}$ , where $SNPs_{9s}$ is the number of SNPs to retain in the nine-spined sticklebacks dataset, $SNPs_{3s}$ is the total number of SNPs passing quality filters in the three-spined sticklebacks dataset, and $L_{3s}$ and $L_{9s}$ are the total number of sequenced sites passing quality control in the three and nine-spined sticklebacks datasets, respectively. This resulted in 1,355,325 SNPs in the nine-spined stickleback dataset (10 1,598,319/129,356,224*1,725,617), which we subsampled from the 1,725,617 SNPs that passed quality filtering. Ultimately, 882,125 SNPs for the three- and 1,355,325 SNPs for the nine-spined stickleback (in form of genotype likelihoods) were used in the downstream comparative LDna analyses.

#### 5 | Testing associations between “synthetic alleles” from LD-clusters and ecotypes using linear mixed models

Since population demographic events such as range expansions, bottlenecks and population growth greatly increase the variance of test statistics from “Genome-scan” based outlier methods, such methods are prone to inflated rates of false positives as well as lack power when background differentiation due to population structuring is high (Galloway et al. 2020; Hoban et al. 2016; Matthey-Doret & Whitlock 2019). Genetic regions underlying variation in locally adapted traits can be identified directly by linkage mapping or genome-wide association (GWA) approaches (e.g. Robinson et al. 2014; Savolainen et al. 2013; Vinkhuyzen et al. 2013). When studying local adaptation, however, both genome-scans and GWA-analyses estimate the same statistical association, namely that between locally adapted phenotypes and genotypes. Since all locally adapted traits are by definition in some way correlated with habitat, the expectation is that all locally adapted traits deviate from the null distribution in GWA-analyses, even if only one of them is tested (LD caused by local adaptation/parallel evolution; Kemppainen et al. 2015). Conversely, if ecotype designation/habitat itself is treated as a binary trait, all genomic regions correlated with local adaptation are expected to deviate from the null distribution. While relatedness in GWA-studies at any level of the population hierarchy (from families to populations) can also lead to false statistical associations between loci and traits of interest (i.e. *p*-value inflation; Kang et al. 2010; Kang et al. 2008), this has elegantly been solved by linear mixed model (LMM) analyses that successfully correct for the genetic relatedness in the data (Kang et al. 2008). When phenotypic values are spatially auto correlated (i.e. certain phenotypic values are mainly shared among related groups of individuals) this can, however, substantially decrease the power in GWA-studies (Kang et al. 2008). The converse can also be true; when two independently colonized and genetically divergent populations share the same heritable phenotypes, correcting for relatedness could potentially increase power to identify the associated loci (Kang et al. 2008). The main benefit of using GWA analyses to find loci associated with local adaptation is thus the ability to use a linear mixed model analysis framework adopted from GWA-studies to monitor and control for relatedness and other co-factors. Indeed, in a related field, Kemppainen et al. 2017 showed that in artificial selection experiments allele frequency change before and after selection also are prone to false positives due to relatedness, and that this could be solved by LMM accounting for relatedness by analysing survival as a binary trait (e.g., “1” for individuals remaining in the population after selection and “0” for those individuals that died). In Kemppainen et al. 2017, the correlation between -log10(*P*)-values from LMM analyses and those attained from a null-distribution generated by permutation was close to unity.

Previously, the Euclidean distance between group centroids (along the two first PC-axes and adjusting for the Euclidean distances within groups), known as the cluster separation score *CSS* has been applied to detect genomic regions associated with marine-freshwater parallelism in three-spined sticklebacks (Jones et al. 2012; Kingman et al. 2020). The significance of *CSS* can be tested by permutation and it correlates with $F_{ST}$, but with higher resolution when differentiation is strong (Jones et al. 2012). Since PC1-coordinates from the LD-clusters typically explain >>90% of the total variation, the *CSS* in this study will be simplified to the difference in means of the PC1-coordinates between marine and freshwater individuals. This simplification, however, is likely to lead to significant loss of information when a window-based approach to *CSS* is applied, i.e. disregarding correlation structure between SNPs, as in e.g. Jones et al. 2012 and Kingman et al. 2020, and is thus only applicable in combination with complexity reduction using e.g. LDna.

Similarly to Kemppainen et al. 2017, where the significance of allele frequency differences between groups of individuals was tested for each bi-allelic SNP separately, we here show that the correlation between -log10(*P*)-values from permutation (performed as in Jones et al. 2012 and Kingman et al. 2020) and from LMM-analyses (treating ecotype as a binary trait, but not correcting for any other potentially confounding factors) also is close to unity (Supplementary Figure 3). Thus, PC1 coordinates from the LD-clusters can be viewed as “multi-locus synthetic alleles” (MLSAs). Here, we thus adopt a similar approach to Kemppainen et al. 2017 to test for MLSA frequency differences between ecotypes using LMM, which allowed us to correct for potential p-value inflation due to relatedness and other potentially confounding factors. Note that in contrast to the *CSS* score used in Jones et al. 2012 and Kingman et al. 2020, our simplified *CSS* scores – that can be replaced by LMM analyses treating ecotype as binary trait – does not account for within group variances. However, the correlation between -log10(*P*)-values from our simplified *CSS* and the *CSS* used in Jones et al. 2012 and Kingman et al. 2020 is high ($r^2$=0.97, n=343; this test was performed on all nine-spined stickleback LDna-1/LDna-2 clusters with parameter settings $|E|_{min}$=20m, $SNP_{min}$=20, with $10^5$ permutation replicates). Similarly, based on all LDna-1 clusters from the three-spined sticklebacks in this study, the linear correlation between -log10(*P*) from LMM and their 95% quantile $F_{ST}$ (based on the loci from each cluster) was $r^2$=0.92 (n=4412). Thus, when not accounting for any confounding factors, testing for the association between MLSA frequency differences between marine and freshwater ecotypes using LMM can replace both *CSS* (as performed in Jones et al. 2012 and Kingman et al. 2020) and $F_{ST}$.

#### 6 | Restricted maximum likelihood (REML) analyses

Linear mixed models can be used to control for hidden population and/or family structure in the data by including the relatedness among individuals as a random effect. Restricted maximum likelihood (REML)-based approaches have been widely used to evaluate the regression parameters and variance components in such models (Kang et al. 2010; Kang et al. 2008). EMMAX (Kang et al. 2010) estimates the variance components once based on an intercept model and subsequently fixes them to evaluate the effect and statistical significance of the SNPs. This approach is preferred over its predecessor due to its computational efficiency while maintaining both power and the ability to control for false positives (Kang et al. 2010; Kang et al. 2008). Association tests here are based on the coordinates of the first PC extracted from PCA (i.e. the SMLAs) on all loci from a given LD-cluster (Li et al. 2018; Fig. 1c) defined as:

$$y_i = \theta_0 + \sum_{l=1}^{m_k} W_{il}\theta_l + u_i + \varepsilon_i, \varepsilon_i \quad N\left(0, \sigma_e^2\right)$$

where  represents the PC coordinates in *k*th LD-cluster; where M is the total number of clusters),  is the intercept, $W_{il}(l=1,\ldots,m_k)$ is the regression parameter of the given PC and is the random effect defined as  with known n x n sized relationship (kinship) matrix **A** and unknown variance  (Li et al. 2018). Given uncertainties in low coverage whole genome sequence data, we obtained the relationship matrix **A** (based on all available loci) in a probabilistic framework from individual allele frequencies estimated iteratively based on inferred population structure from genotype likelihoods using the software *PCAngsd* (Korneliussen et al. 2014).

#### 7 | Correction for multiple testing by permutation

Although computationally much more intensive, an alternative but potentially less conservative alternative to the “false discovery rate” method to control the family-wise error (FWER) is by permutation that accounts for the correlation structure among the multiple tests (Efron 2010; Joo et al. 2016; Li et al. 2018). This method is particularly useful in combination with LD-based complexity reduction, where the total number of tests can in many cases substantially be lowered, as demonstrated by Li et al. 2018. To correctly account for the among individual correlation structure in the data we generated phenotypic data under the null-hypothesis by drawing phenotypic data from a multivariate normal distribution $\theta_0$ and then using EMMAX to calculate the test-statistics on each sample. The resulting empirical distributions of the test statistics were then used to adjust p-values to control the multiplicity as in Li et al. 2018. Since we are testing for associations between loci and ecotype (treated as a binary trait), we treated the phenotypes as threshold traits, with the number of freshwater and marine ecotypes equal to that observed in the real data.

#### 8 | Genetic diversity and divergence in the marine–freshwater divergent genomic regions

To see whether genomic regions under selection have more ancient origins than neutral regions, we tested whether the absolute genetic divergence ($d_{XY}$, Nei 1987) in the genomic regions under selection exceeds that in the rest of the genome. All freshwater populations considered are the results of postglacial invasions, and hence mean $d_{XY}$ between freshwater and geographically close marine populations is expected to be low, i.e. to approximate $\pi$ from the ancestral population (see discussion in Ravinet et al. 2017). If selection is acting on a novel or rare variant, we can expect a high peak in $F_{ST}$ but not an increase in $d_{XY}$, as allelic differentiation would be a result of reduced $\pi$ in the population experiencing selection (i.e. a selective sweep; Cruickshank & Hahn 2014). In contrast, if selection is acting on ancient haplotypes, a peak in $F_{ST}$ is likely to be driven by an increase in $d_{XY}$, rather than a decrease in $\pi$ (e.g. Nelson & Cresko 2018). Comparing $d_{XY}$ in regions underlying parallel evolution with the rest of the genome could provide information on the evolutionary origin of the SGV under selection, but such comparison is not straightforward. Like other measures of diversity ($\pi$ and $\theta$), $d_{XY}$ is strongly affected by the heterogeneity in background selection, recombination rates and mutation rates across the genome (Charlesworth et al. 1993). Since the processes that govern diversity levels within genomes (background selection, mutation rate and recombination rate variation) are conserved between closely related populations (and species), different measures of diversity $\pi$ and $d_{XY}$) are correlated across the genome of closely related populations. Dutoit et al. 2017 demonstrated this correlation is maintained even after lineage sorting is complete, suggesting shared ancestral variation cannot explain this pattern.

Here we take advantage of this correlation to detect whether genomic regions under selection show excess absolute divergence. We start from the assumption that marine populations within the same regions are proxies of ancestral diversity. For each genomic region involved in parallel evolution we then run a linear model where $\pi_{marine}$ is the predictor of $d_{XY}$ across the genome, including 100 random genomic regions as well as the region under selection. This yields a distribution of residuals ($\Delta d_{XY}$), where a $\Delta d_{XY}$ of 0 represents the $d_{XY}$ predicted by the ancestral diversity. This is conceptually similar to Nei's measure of relative divergence — $d_a = d_{XY} - (\pi_x\ \pi_y)/2$, (Nei & Li 1979) — but with the difference that we use $\pi$ *p* of the marine population as a proxy of ancestral diversity rather than the mean of $\pi$ between both populations. Furthermore, the residuals are cantered so that positive and negative values represent higher and lower relative divergence than the mean across all regions. We use the random 100 neutral genomic regions to infer the confidence intervals for the neutral distribution, and then compare to $\Delta d_{XY}$ of the target region. We use a similar approach to detect whether regions under selection show lower diversity in the freshwater populations than the rest of the genome (i.e. if peaks in $F_{ST}$ are caused by selective sweeps). As for $d_{XY}$, we run a linear regression between $\pi_{freshwater}$ and $\pi_{marine}$ (lm $\pi_{freshwater} \sim \pi_{marine}$) based on the same 101 genomic regions. We refer to the residuals of the model as $\Delta\pi$. A negative $\Delta\pi$ in the genomic region under selection would be suggestive of a reduction in $p$ relative to the expected difference in $p$ between the two species.

Estimates of genetic diversity ($\pi$ and $d_{XY}$) were calculated as follows. Focusing on the Norwegian Sea freshwater three-spined sticklebacks and White & Barents Seas freshwater nine-spined sticklebacks (see Materials and Methods), we treated one focal freshwater population and one marine population across the Atlantic as a marine–freshwater population pair. Only the populations with at least two samples were considered and all population pairs were found. For each candidate genomic region in a LD-cluster (5kbp upstream plus 5kbp downstream of the loci with the highest $F_{ST}$), we calculated $p$ in each ecotype and $d_{XY}$ between the two ecotypes in each population pair. We first estimated 1D-SFS of each population and 2D-SFS of both populations using ANGSD. Then we calculated $\pi$ and $d_{XY}$ based on 1D-SFS and 2D-SFS respectively using custom scripts derived from Momigliano et al. 2021. Scripts can be found in: github.com/Nopaoli/Demographic-Modelling/tree/master/Diversity_fromSFS**.** Models of $\Delta d_{XY}$ *and* $\Delta\pi$ were performed in R for each population pair, and the results ($\Delta d_{XY}$, $\Delta\pi$ and $r^2$) were averaged across all population pairs in Fig. 5 and Supplementary Fig. 5 for presentation.

**Supplementary Figure 1 | Sampling maps of the populations used in different analyses.** (a) All populations in the study. (b) Populations used in the analyses of individual heterozygosity (*H*). (c) Populations used in the analyses of differentiation ($F_{ST}$), nucleotide diversity (*p*) and Watterson's theta ($\theta$). (d) European populations used in IBD analyses. (e) Populations used in SNAPP analyses. Population identities in Fig. 3 are given on the bottom-right. See Method for justifications for the sampling in different analyses.

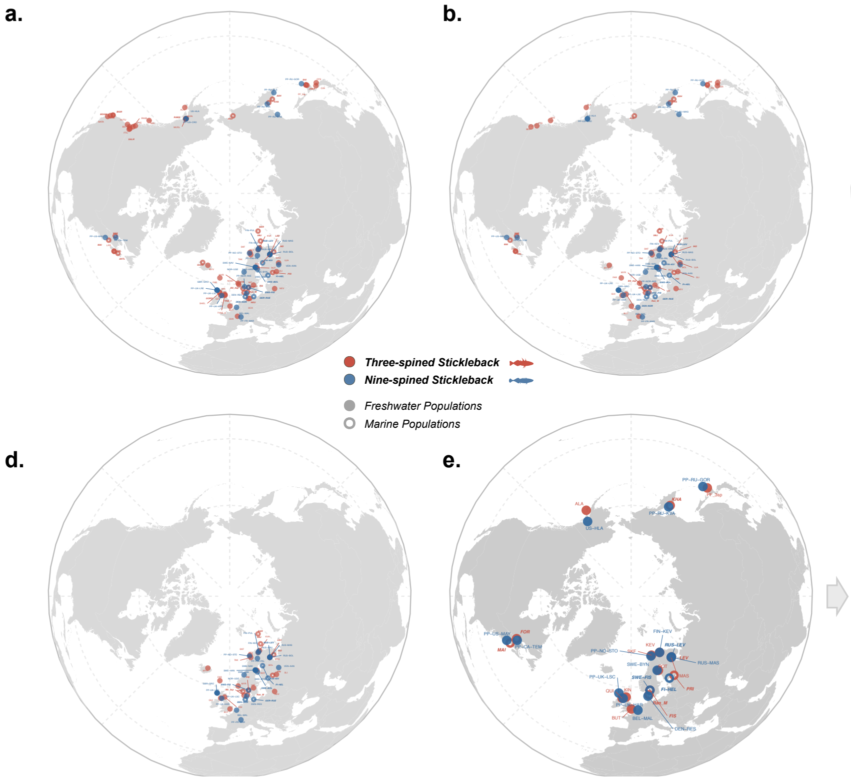


**Populations in the Comparative SNAPP Analyses**

Three-spined stickleback

| Population ID | Population ID in the Fig. 3 |
|---|---|
| MAI | MAI |
| FOR | FOR |
| MAS | MAS |
| LEV | LEV |
| BUT | BUT |
| SKF | SKF |
| ROT | ROT |
| KEV | KEV |
| KIN | KIN |
| QUI | QUI |
| Ran_M | Ran |
| PRI | PRI |
| FIS | FIS |
| PF_3sp | PF |
| KHA | KHA |
| ALA | ALA |

Nine-spined stickleback

| Population ID | Population ID in the Fig. 3 |
|---|---|
| US-HLA | HLA |
| PP-RU-KVA | KVA |
| PP-RU-GOR | GOR |
| PP-CA-TEM | TEM |
| PP-US-MAY | MAY |
| PP-UK-HAR | HAR |
| PP-UK-LSC | LSC |
| BEL-MAL | MAL |
| SWE-FIS | FIS |
| DEN-RES | RES |
| PP-NO-STO | STO |
| FIN-KEV | KEV |
| RUS-LEV | LEV |
| RUS-MAS | MAS |
| SWE-BYN | BYN |
| FI-HEL | HEL |

**Supplementary Figure 2 | Nested three-step LDna-complexity reduction.** First, (a) sets of highly correlated loci (as indicated by heatmap where light shade indicates stronger LD) along windows within chromosomes can be grouped together into LD-clusters, despite them being interspersed and overlapping along the chromosome due to "mosaic-like" LD. From each cluster one rSNP (connected to all other SNPs in its cluster by high LD) is taken to represent the whole cluster in LDna-2. Second, (b) using an LD-matrix $K_{LD}$ between all rSNPs from (a) and single linkage clustering standard LDna is performed separately for each chromosome. Here, each branch is considered as an LD-cluster, with the size and correlation determined by $|E|_{min}$ (the minimum number of edges in the cluster; higher values yielding larger but less correlated LD-clusters). Third, (c), a correlation matrix $K_{PC1}$ was constructed based on the squared correlation coefficients between MLSAs for all pairwise comparisons between LDna-1 clusters which is then used to construct a genome wide correlation tree. Note that in LDna-3 all LDna-2 clusters, regardless of chromosome, are included. All LD-clusters identified in LDna-1 that are not part of any LDna-2 clusters but contain $>SNP_{min}$ number of loci are also included in LDna-3 analyses. Lastly, any LDna-1/2 clusters correlated at $>Cor_{th}$ will be considered a single independent LDna-3 cluster. This step can significantly reduce the number of tests by pooling together genome wide loci that only reflect population structuring.

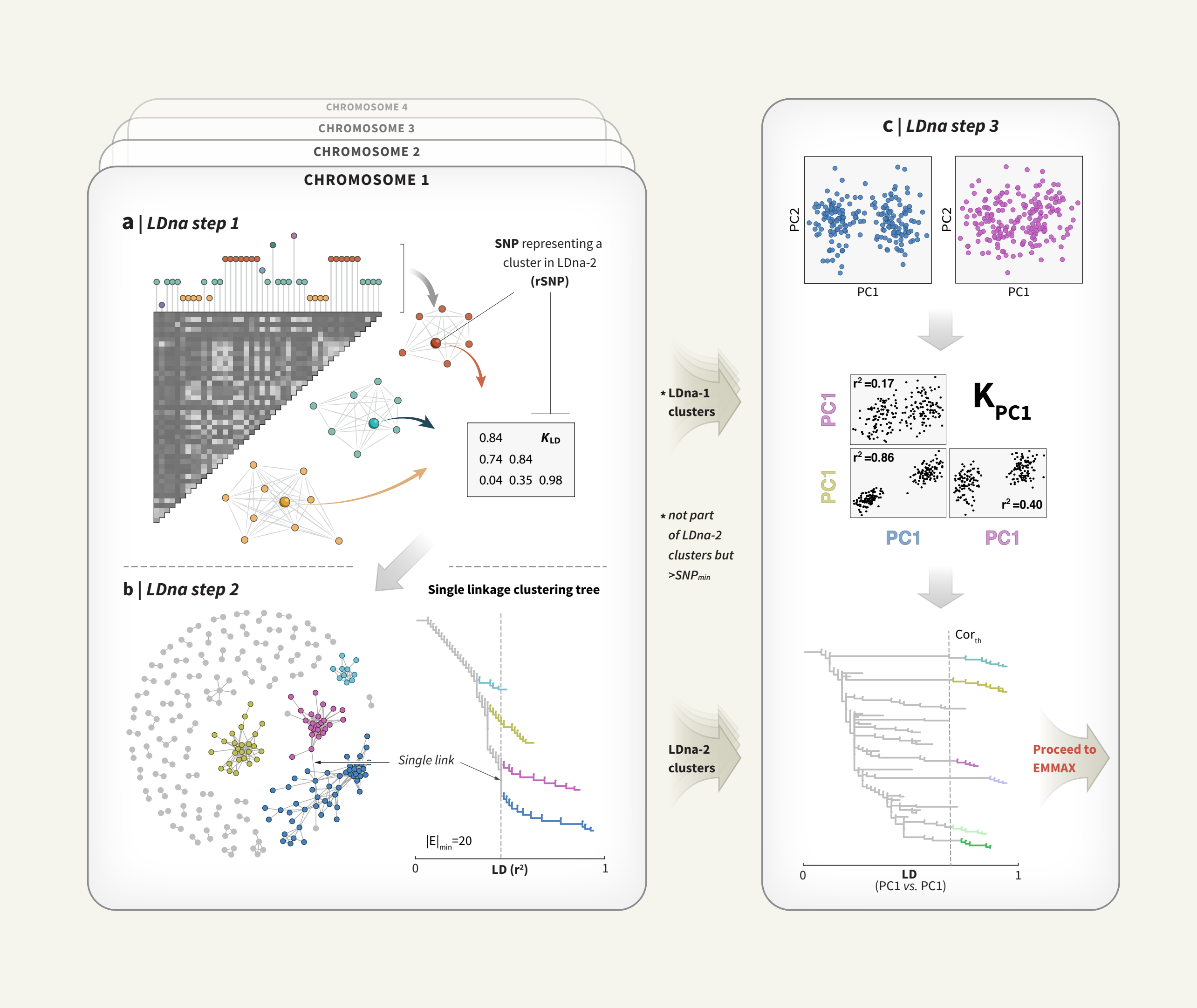

**Supplementary Figure 3 | The relationship between EMMAX analyses performed on "synthetic alleles" from PCA-analyses of LD-clusters.** Data points represent analyses from 1000 randomly chosen LDna-1 clusters from the three- and nine spined sticklebacks data sets testing for associations between SMLAs (PC1-coordinates from LD-clusters) and ecotype either by (y-axis) EMMAX, treating ecotype as a binary trait and not including any co-factors or random effects or by (x-axis) permutation (n=$10^5$). In both species, the relationship is close to unity, with slightly larger variance for nine-spined sticklebacks.

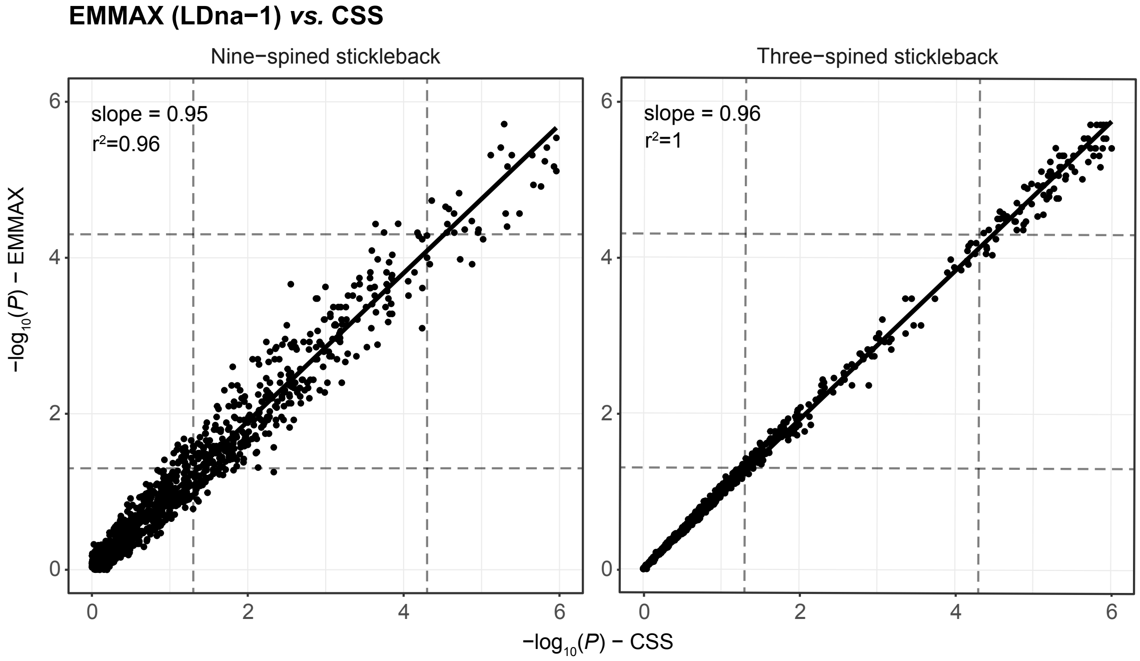

**Supplementary Figure 4 | Cumulative numbers of outlier regions detected for randomly selected parameter/correction method combinations.** Genomic regions associated with marine-freshwater differentiation was tested with 45 orthogonal parameter combinations of $|E|_{min}$, $SNP_{min}$ and $Cor_{th}$ and four methods to correct for multiple testing and p-value inflation ("A+fdr, "A+perm", "GC+fdr" and "perm", se main text for details). Black lines show the mean (n=100) cumulative increase of significant outlier regions when sequentially including more parameter/correlation method combinations (in total 180) in random order, with solid line indicating regions with $C \geq 0$, dashed line indicating regions with $C \geq 0.05$ and dotted line indicating regions with $C \geq 0.1$. In each panel from left to right the level of one parameter is fixed while other parameter combinations are randomly sampled. This shows the influence of each parameter on the number of outlier regions. For instance, the most "successful" $|E|_{min}$ settings were $|E|_{min}$=20 for three-spined but $|E|_{min}$=10 for nine-spined sticklebacks (left panel). Similarly, the most successful $Cor_{th}$ value for three-spined sticklebacks was $Cor_{th}$=10, while this parameter had less influence for nine-spined sticklebacks (right panel). Both reflect the fact that outlier regions affect a larger number of correlated loci in three- compared to nine-spined sticklebacks and that a given parameter setting can be more successful (in detecting significant outlier regions) in one species compared to another.

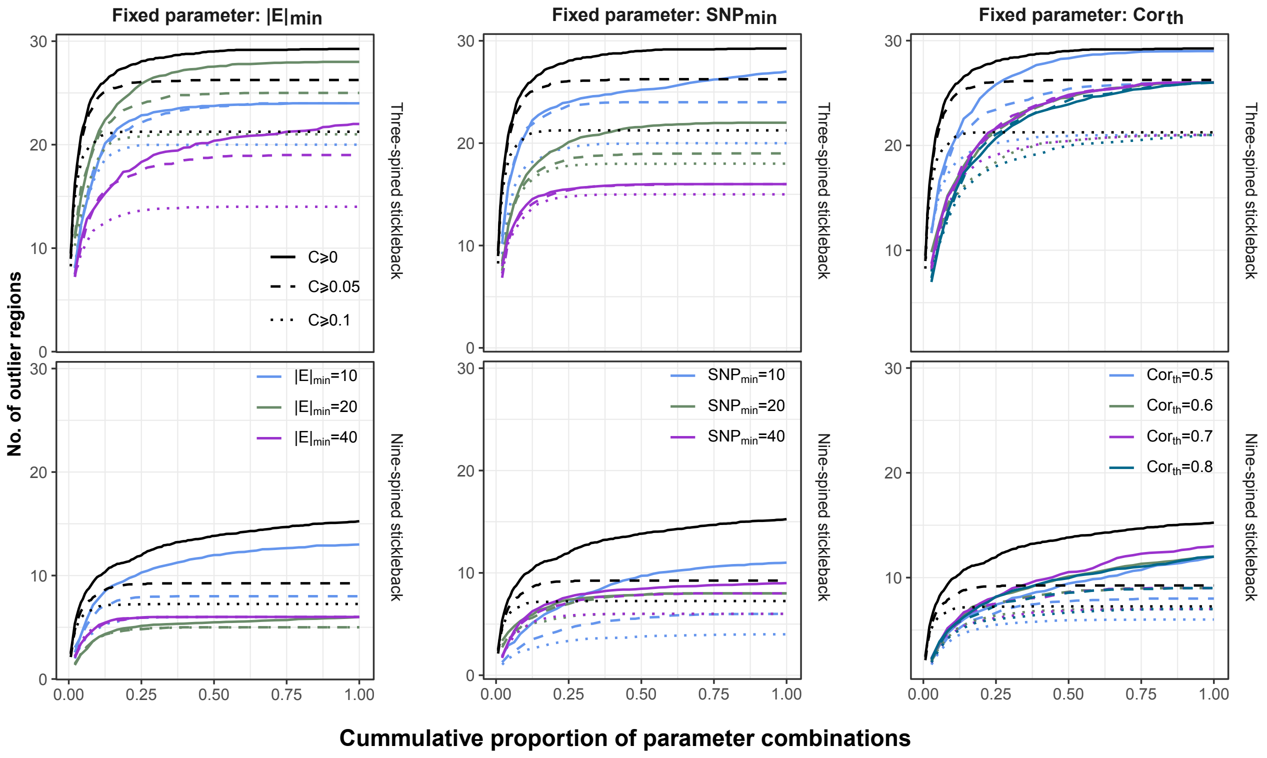

**Supplementary Figure 5 | The maximum-clade-credibility summary trees of the SNAPP phylogenies reported in Fig. 3.** (a) The tree of nine-spined sticklebacks. (b) The tree of three-spined sticklebacks. The numbers in the nodes represent divergence time estimates in form of “mean divergence age (Mya) [range of 95% highest posterior density (HPD) intervals]”. The circles (A, B and C) indicate the calibration points applied in the SNAPP analyses. The populations in red rectangles are used to analyse the correlation between divergence times and parallelism (see Materials and methods for details and population abbreviations).

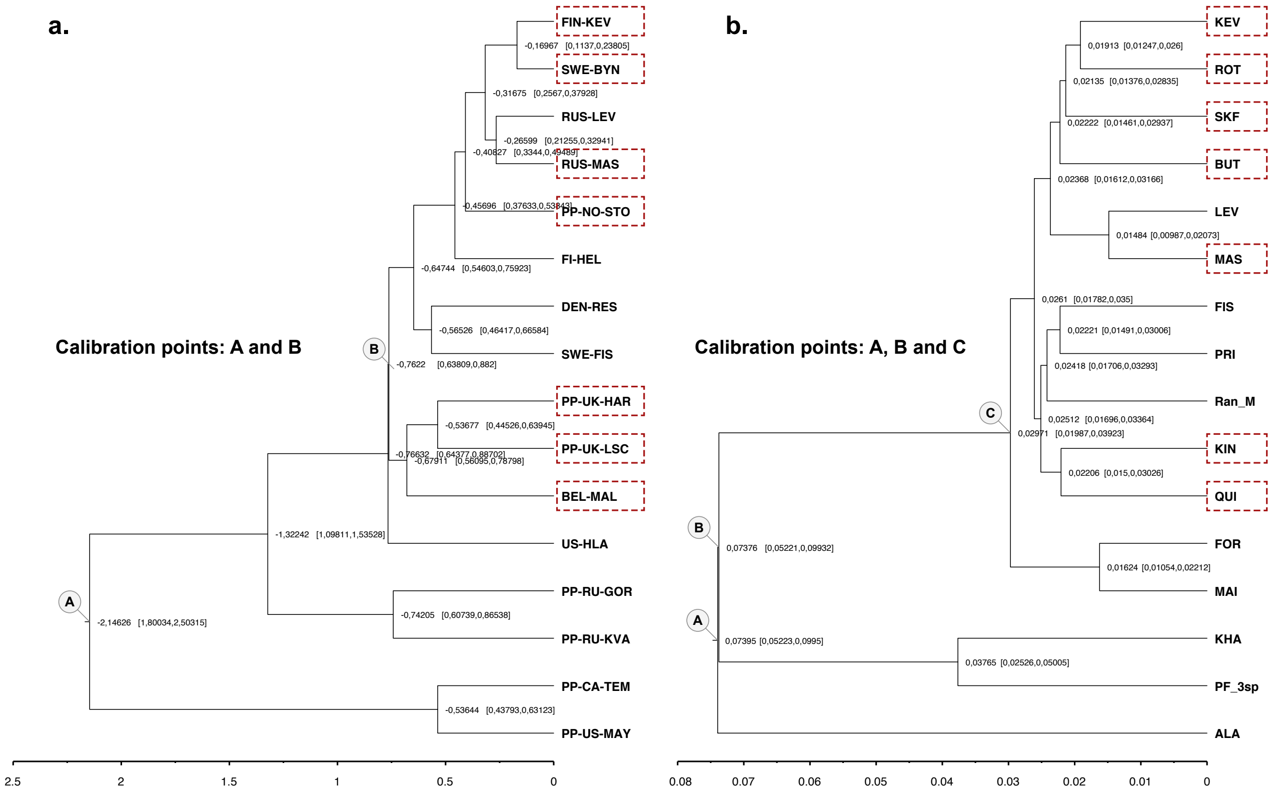

**Supplementary Figure 5 | Supplementary genetic diversity estimates.** Genetic diversity between the two species: (a, b) Watterson's theta (*θ*), (c) nucleotide diversity (*π*) and (d) heterozygosity (*H*). (b-d) Genetic diversity incorporating the known hybridised nine-spined stickleback populations in analyses (see Materials and methods). (e) The *H* between admixed populations and non-admixed populations of nine-spined sticklebacks. Statistics are reported in the results.

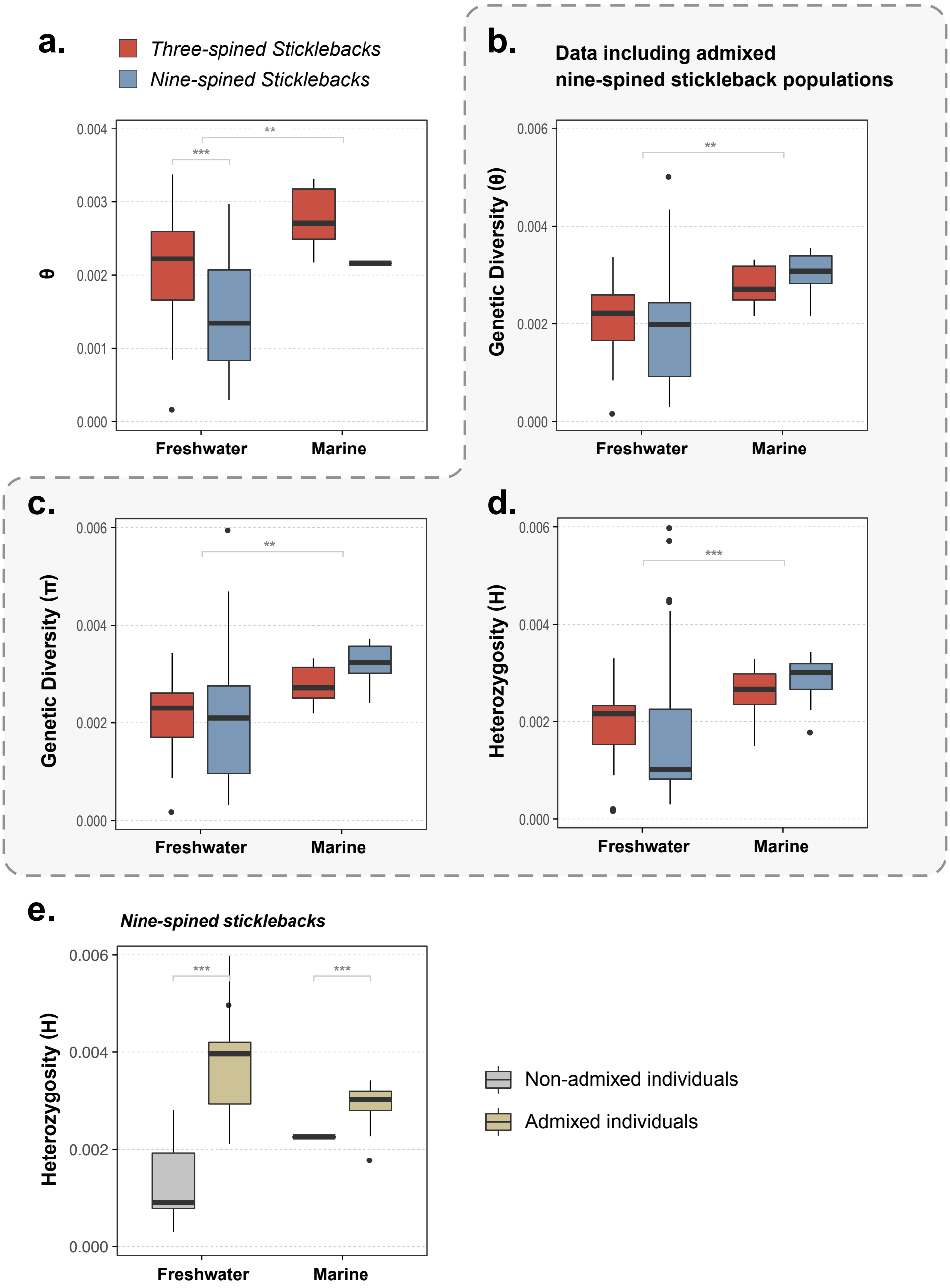

**Supplementary Figure 7 | Intra-clade genetic differentiation.** Boxplots of pairwise $F_{ST}$ between populations within genetic clades in nine-spined sticklebacks compared to three-spined sticklebacks. Genetic differentiation is higher in nine-spined sticklebacks whether we consider freshwater-freshwater, marine-freshwater or marine-marine comparisons. Note that data for all ecotype contrasts were not available for all clades.

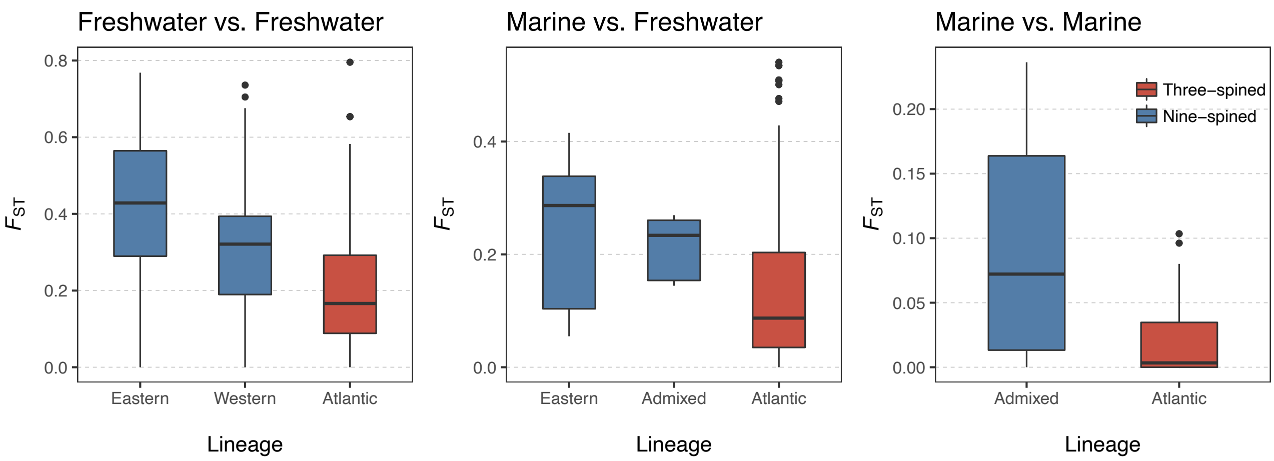

**Supplementary Figure 8 | Genetic diversity and divergence in the genomic regions under selection.** Data for three- (No. 1-26) and nine-spined sticklebacks (No. 27-35) are presented. In each panel, the upper plot indicates the position of the candidate region with red arrow. The lower plot summarises the residuals of linear regression models based on the genetic diversity ($\Delta\pi$) and genetic divergence ($\Delta d_{XY}$) derived from marine–freshwater population pairs in corresponding LD-cluster (see Materials and methods). The correlation coefficient ($R^2$) is shown as an averaged value across all models from different population pairs. All models are statistically significant ($P$<0.001).

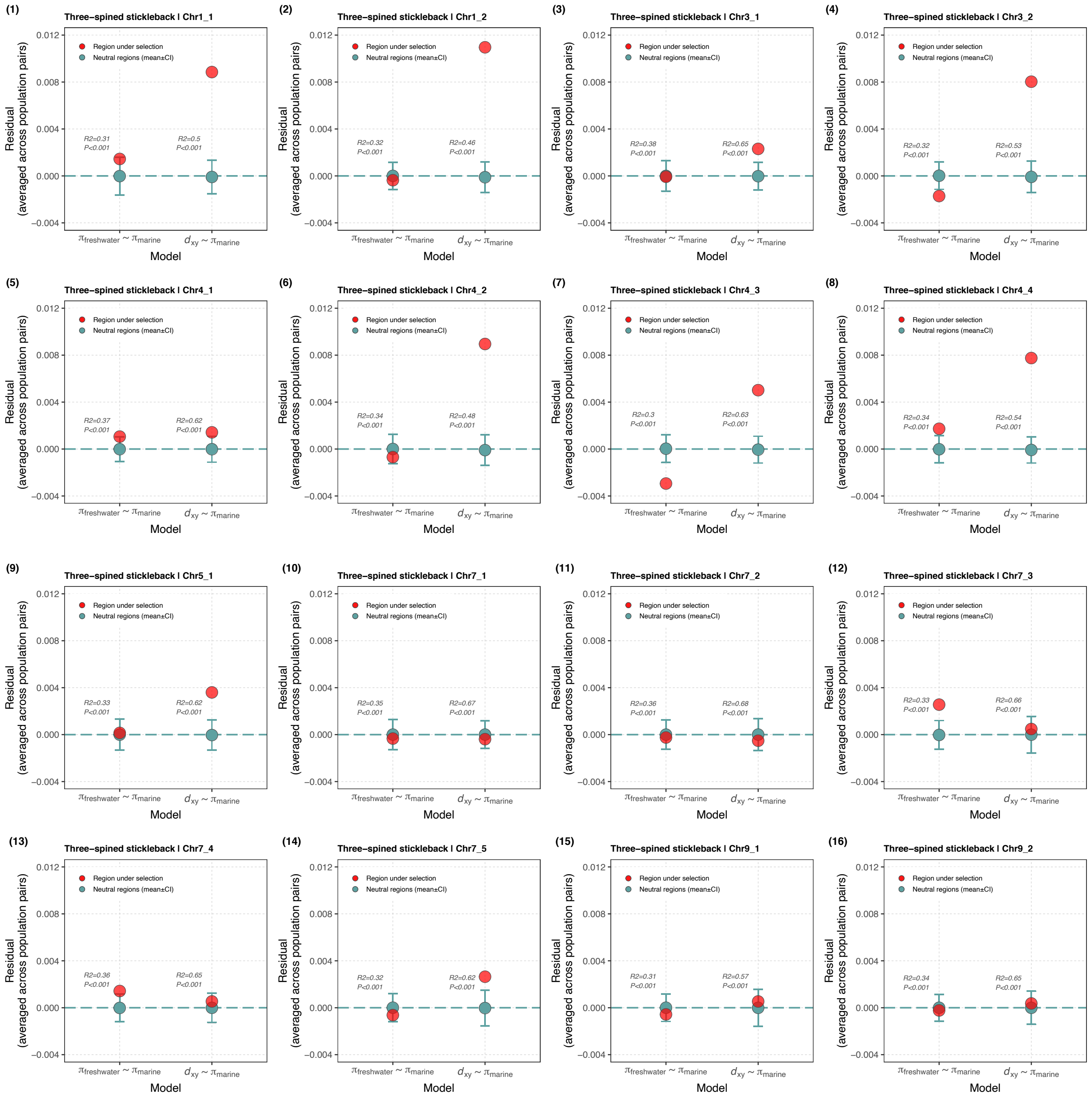

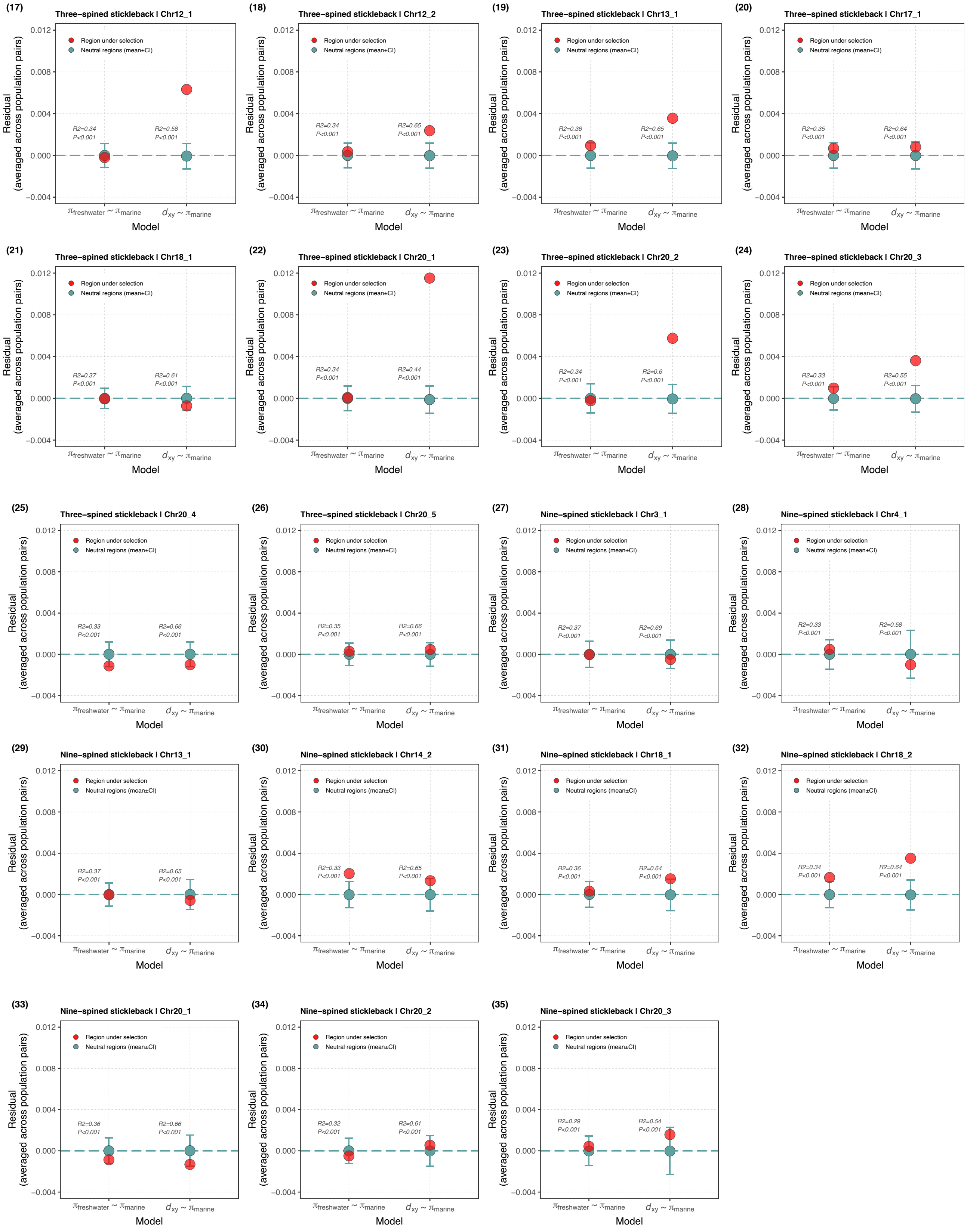

Residual (averaged across population pairs)
Model
Region under selection
Neutral regions (mean±CI)
πfreshwater ~ πmarine
dxy ~ πmarine
(17) Three−spined stickleback | Chr12_1
R2=0.34 P<0.001
R2=0.58 P<0.001
(18) Three−spined stickleback | Chr12_2
R2=0.34 P<0.001
R2=0.65 P<0.001
(19) Three−spined stickleback | Chr13_1
R2=0.36 P<0.001
R2=0.65 P<0.001
(20) Three−spined stickleback | Chr17_1
R2=0.35 P<0.001
R2=0.64 P<0.001
(21) Three−spined stickleback | Chr18_1
R2=0.37 P<0.001
R2=0.61 P<0.001
(22) Three−spined stickleback | Chr20_1
R2=0.34 P<0.001
R2=0.44 P<0.001
(23) Three−spined stickleback | Chr20_2
R2=0.34 P<0.001
R2=0.6 P<0.001
(24) Three−spined stickleback | Chr20_3
R2=0.33 P<0.001
R2=0.55 P<0.001
(25) Three−spined stickleback | Chr20_4
R2=0.33 P<0.001
R2=0.66 P<0.001
(26) Three−spined stickleback | Chr20_5
R2=0.35 P<0.001
R2=0.66 P<0.001
(27) Nine−spined stickleback | Chr3_1
R2=0.37 P<0.001
R2=0.69 P<0.001
(28) Nine−spined stickleback | Chr4_1
R2=0.33 P<0.001
R2=0.58 P<0.001
(29) Nine−spined stickleback | Chr13_1
R2=0.37 P<0.001
R2=0.65 P<0.001
(30) Nine−spined stickleback | Chr14_2
R2=0.33 P<0.001
R2=0.65 P<0.001
(31) Nine−spined stickleback | Chr18_1
R2=0.36 P<0.001
R2=0.64 P<0.001
(32) Nine−spined stickleback | Chr18_2
R2=0.34 P<0.001
R2=0.64 P<0.001
(33) Nine−spined stickleback | Chr20_1
R2=0.36 P<0.001
R2=0.66 P<0.001
(34) Nine−spined stickleback | Chr20_2
R2=0.32 P<0.001
R2=0.61 P<0.001
(35) Nine−spined stickleback | Chr20_3
R2=0.29 P<0.001
R2=0.54 P<0.001

**Supplementary Figure 9 | Correlations between divergence time and genetic parallelism.** The correlation between divergence time and genetic parallelism was tested in (a) three- and (b) nine-spined stickleback population pairs, respectively. Seven pairs of populations shown in the phylogenies in Fig. ٣ were used in these analyses. Specific populations are presented in the Supplementary Fig. ٣. Each dot represents a pairwise comparison. The x-axis presents the divergence time (Mya) of the populations in each pair, while the y-axis displays the proportion of loci involved showing genetic parallelism in each population pair. The correlation between divergence times and genetic parallelism was tested using maximum-likelihood population-effects (MLPE) model.

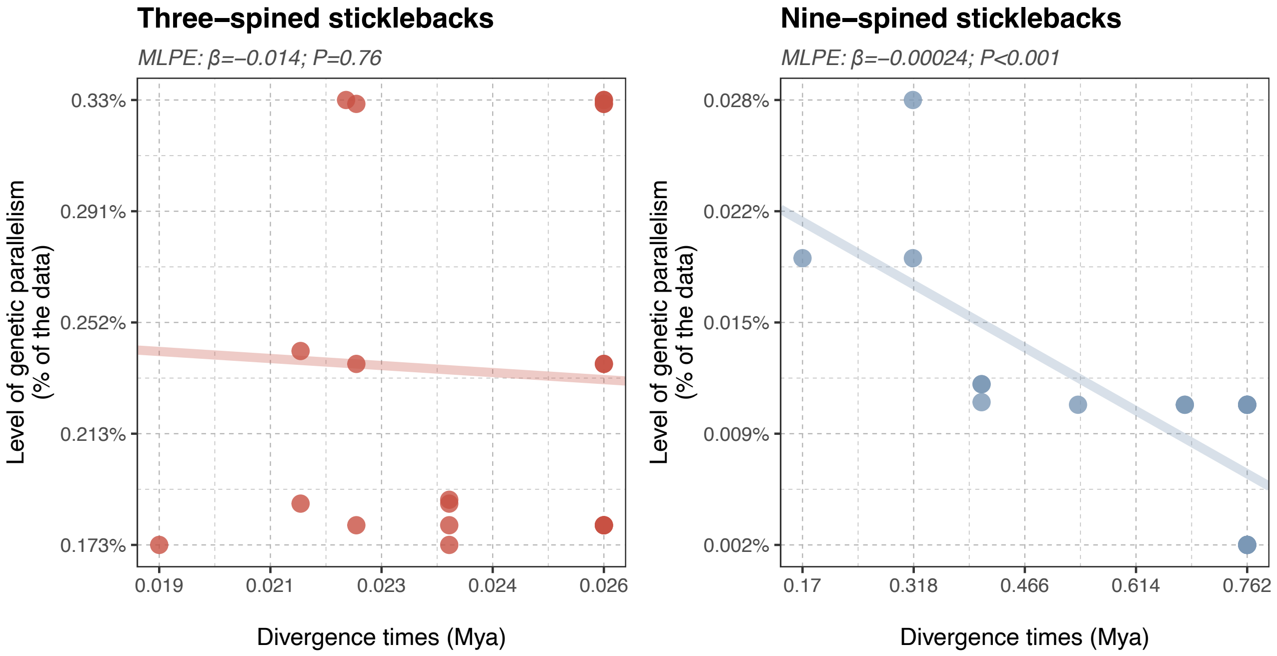

**Supplementary Table 2** | Information on the nine-spined stickleback samples used in the study. The information on the three-spined stickleback samples is available at the Supplementary Table 1 in Fang et al. (2020).

| Sample_ID | Population_ID | Ecotype | Region | GPS_N | GPS_E | Coverage |
|---|---|---|---|---|---|---|
| RUS-BOL-11 | RUS-BOL | Freshwater | White and Barents Seas | 66.3 | 33.4 | 17.59 |
| RUS-BOL-12 | RUS-BOL | Freshwater | White and Barents Seas | 66.3 | 33.4 | 13.543 |
| RUS-BOL-13 | RUS-BOL | Freshwater | White and Barents Seas | 66.3 | 33.4 | 15.704 |
| RUS-BOL-6 | RUS-BOL | Freshwater | White and Barents Seas | 66.3 | 33.4 | 14.357 |
| RUS-BOL-8 | RUS-BOL | Freshwater | White and Barents Seas | 66.3 | 33.4 | 15.359 |
| RUS-BOL-9 | RUS-BOL | Freshwater | White and Barents Seas | 66.3 | 33.4 | 18.188 |
| RUS-MAS-4 | RUS-MAS | Freshwater | White and Barents Seas | 66.3 | 33.4 | 10.793 |
| RUS-MAS-5 | RUS-MAS | Freshwater | White and Barents Seas | 66.3 | 33.4 | 22.259 |
| RUS-MAS-6 | RUS-MAS | Freshwater | White and Barents Seas | 66.3 | 33.4 | 11.077 |
| RUS-MAS-7 | RUS-MAS | Freshwater | White and Barents Seas | 66.3 | 33.4 | 10.008 |
| RUS-MAS-8 | RUS-MAS | Freshwater | White and Barents Seas | 66.3 | 33.4 | 10.732 |
| RUS-MAS-9 | RUS-MAS | Freshwater | White and Barents Seas | 66.3 | 33.4 | 10.241 |
| RUS-KRU-1 | RUS-KRU | Freshwater | White and Barents Seas | 66.3 | 33.4 | 10.448 |
| RUS-KRU-11 | RUS-KRU | Freshwater | White and Barents Seas | 66.3 | 33.4 | 10.047 |
| RUS-KRU-3 | RUS-KRU | Freshwater | White and Barents Seas | 66.3 | 33.4 | 10.287 |
| RUS-KRU-4 | RUS-KRU | Freshwater | White and Barents Seas | 66.3 | 33.4 | 14.49 |
| RUS-KRU-5 | RUS-KRU | Freshwater | White and Barents Seas | 66.3 | 33.4 | 34.787 |
| RUS-KRU-9 | RUS-KRU | Freshwater | White and Barents Seas | 66.3 | 33.4 | 13.176 |
| FIN-PUL-1 | FIN-PUL | Freshwater | White and Barents Seas | 69.967 | 27.967 | 16.138 |
| FIN-PUL-2 | FIN-PUL | Freshwater | White and Barents Seas | 69.967 | 27.967 | 17.165 |
| FIN-PUL-3 | FIN-PUL | Freshwater | White and Barents Seas | 69.967 | 27.967 | 18.294 |
| FIN-PUL-4 | FIN-PUL | Freshwater | White and Barents Seas | 69.967 | 27.967 | 15.586 |
| FIN-PUL-5 | FIN-PUL | Freshwater | White and Barents Seas | 69.967 | 27.967 | 14.841 |
| FIN-PUL-6 | FIN-PUL | Freshwater | White and Barents Seas | 69.967 | 27.967 | 14.959 |
| FIN-KEV-1 | FIN-KEV | Freshwater | White and Barents Seas | 69.75 | 27.017 | 14.231 |
| FIN-KEV-2 | FIN-KEV | Freshwater | White and Barents Seas | 69.75 | 27.017 | 16.671 |
| FIN-KEV-3 | FIN-KEV | Freshwater | White and Barents Seas | 69.75 | 27.017 | 17.047 |
| FIN-KEV-4 | FIN-KEV | Freshwater | White and Barents Seas | 69.75 | 27.017 | 17.031 |
| FIN-KEV-5 | FIN-KEV | Freshwater | White and Barents Seas | 69.75 | 27.017 | 14.689 |
| FIN-KEV-8 | FIN-KEV | Freshwater | White and Barents Seas | 69.75 | 27.017 | 14.374 |
| PP-VEN-AAN-2 | VEN-AAN | Freshwater | White and Barents Seas | 61.583 | 34.633 | 12.064 |
| PP-VEN-AAN-8 | VEN-AAN | Freshwater | White and Barents Seas | 61.583 | 34.633 | 8.887 |
| SWE-ABB-411 | SWE-ABB | Freshwater | Baltic Sea | 64.478 | 19.436 | 9.954 |
| SWE-ABB-424 | SWE-ABB | Freshwater | Baltic Sea | 64.478 | 19.436 | 10.525 |
| SWE-ABB-45 | SWE-ABB | Freshwater | Baltic Sea | 64.478 | 19.436 | 10.937 |
| SWE-ABB-6 | SWE-ABB | Freshwater | Baltic Sea | 64.478 | 19.436 | 10.977 |
| SWE-BYN-10 | SWE-BYN | Freshwater | Baltic Sea | 64.455 | 19.444 | 11.325 |
| SWE-BYN-11 | SWE-BYN | Freshwater | Baltic Sea | 64.455 | 19.444 | 10.904 |
| SWE-BYN-8 | SWE-BYN | Freshwater | Baltic Sea | 64.455 | 19.444 | 11.38 |
| SWE-NAV-1 | SWE-NAV | Freshwater | Baltic Sea | 64.565 | 19.199 | 16.198 |
| SWE-NAV-4 | SWE-NAV | Freshwater | Baltic Sea | 64.565 | 19.199 | 13.638 |
| SWE-NAV-8 | SWE-NAV | Freshwater | Baltic Sea | 64.565 | 19.199 | 18.135 |
| SWE-HAN-3 | SWE-HAN | Freshwater | Baltic Sea | 64.557 | 19.174 | 10.438 |
| SWE-HAN-4 | SWE-HAN | Freshwater | Baltic Sea | 64.557 | 19.174 | 10.093 |
| SWE-HAN-5 | SWE-HAN | Freshwater | Baltic Sea | 64.557 | 19.174 | 9.695 |
| DEN-RES-1 | DEN-RES | Freshwater | North Sea | 56.183 | 9.633 | 12.722 |
| DEN-RES-4 | DEN-RES | Freshwater | North Sea | 56.183 | 9.633 | 12.617 |
| DEN-RES-6 | DEN-RES | Freshwater | North Sea | 56.183 | 9.633 | 12.551 |
| PP-FIN-PAL-1 | PP-FIN-PAL | Freshwater | White and Barents Seas | 68.029 | 24.157 | 7.002 |
| PP-FIN-PAL-2 | PP-FIN-PAL | Freshwater | White and Barents Seas | 68.029 | 24.157 | 6.851 |
| PP-FIN-PAL-3 | PP-FIN-PAL | Freshwater | White and Barents Seas | 68.029 | 24.157 | 8.823 |
| PP-FIN-PAL-4 | PP-FIN-PAL | Freshwater | White and Barents Seas | 68.029 | 24.157 | 4.618 |
| PP-FIN-PAL-5 | PP-FIN-PAL | Freshwater | White and Barents Seas | 68.029 | 24.157 | 12.016 |
| PP-FIN-PAL-6 | PP-FIN-PAL | Freshwater | White and Barents Seas | 68.029 | 24.157 | 16.961 |
| PHY-NOR-STO-01 | PP-NO-STO | Freshwater | Norwegian Sea | 69.748 | 18.415 | 6.985 |
| PHY-NOR-STO-02 | PP-NO-STO | Freshwater | Norwegian Sea | 69.748 | 18.415 | 7.84 |
| PHY_NOR-STO-261 | PP-NO-STO | Freshwater | Norwegian Sea | 69.748 | 18.415 | 7.773 |
| NOR-UGE-10 | NOR-UGE | Freshwater | Norwegian Sea | 63.958 | 10.427 | 16.52 |
| NOR-UGE-11 | NOR-UGE | Freshwater | Norwegian Sea | 63.958 | 10.427 | 18.62 |
| NOR-UGE-12 | NOR-UGE | Freshwater | Norwegian Sea | 63.958 | 10.427 | 17.442 |
| NOR-UGE-13 | NOR-UGE | Freshwater | Norwegian Sea | 63.958 | 10.427 | 17.418 |
| NOR-UGE-14 | NOR-UGE | Freshwater | Norwegian Sea | 63.958 | 10.427 | 17.632 |
| NOR-UGE-15 | NOR-UGE | Freshwater | Norwegian Sea | 63.958 | 10.427 | 16.113 |
| NOR-UGE-18 | NOR-UGE | Freshwater | Norwegian Sea | 63.958 | 10.427 | 16.045 |
| NOR-UGE-1 | NOR-UGE | Freshwater | Norwegian Sea | 63.958 | 10.427 | 18.116 |
| NOR-UGE-20 | NOR-UGE | Freshwater | Norwegian Sea | 63.958 | 10.427 | 17.078 |
| NOR-UGE-21 | NOR-UGE | Freshwater | Norwegian Sea | 63.958 | 10.427 | 19.514 |
| NOR-UGE-23 | NOR-UGE | Freshwater | Norwegian Sea | 63.958 | 10.427 | 20.286 |
| NOR-UGE-24 | NOR-UGE | Freshwater | Norwegian Sea | 63.958 | 10.427 | 16.662 |
| NOR-UGE-26 | NOR-UGE | Freshwater | Norwegian Sea | 63.958 | 10.427 | 17.674 |
| NOR-UGE-27 | NOR-UGE | Freshwater | Norwegian Sea | 63.958 | 10.427 | 19.212 |
| NOR-UGE-2 | NOR-UGE | Freshwater | Norwegian Sea | 63.958 | 10.427 | 16.919 |
| NOR-UGE-5 | NOR-UGE | Freshwater | Norwegian Sea | 63.958 | 10.427 | 16.361 |
| NOR-UGE-6 | NOR-UGE | Freshwater | Norwegian Sea | 63.958 | 10.427 | 16.295 |
| NOR-UGE-7 | NOR-UGE | Freshwater | Norwegian Sea | 63.958 | 10.427 | 23.668 |
| NOR-UGE-8 | NOR-UGE | Freshwater | Norwegian Sea | 63.958 | 10.427 | 20.279 |
| NOR-UGE-9 | NOR-UGE | Freshwater | Norwegian Sea | 63.958 | 10.427 | 16.924 |
| PHY-FRA-MAR-01 | PP-FR-MAR | Freshwater | Western Europe | 47.024 | 5.254 | 7.771 |
| PHY-FRA-MAR-02 | PP-FR-MAR | Freshwater | Western Europe | 47.024 | 5.254 | 8.37 |
| PP-NOR-AVE-2a | PP-NOR-AVE | Freshwater | North Sea | 58.714 | 9.116 | 5.626 |
| BEL-MAL-F26 | BEL-MAL | Freshwater | Western Europe | 51.175 | 3.469 | 12.671 |
| BEL-MAL-F27 | BEL-MAL | Freshwater | Western Europe | 51.175 | 3.469 | 12.711 |
| BEL-MAL-F28 | BEL-MAL | Freshwater | Western Europe | 51.175 | 3.469 | 11.554 |
| BEL-MAL-F29 | BEL-MAL | Freshwater | Western Europe | 51.175 | 3.469 | 12.355 |
| BEL-MAL-F30 | BEL-MAL | Freshwater | Western Europe | 51.175 | 3.469 | 12.33 |
| PHY-SCO-HAR-01 | PP-UK-HAR | Freshwater | Western Europe | 55.754 | -4.429 | 7.318 |
| PHY-SCO-HAR-02 | PP-UK-HAR | Freshwater | Western Europe | 55.754 | -4.429 | 8.785 |
| PP-UK-LRE-1 | PP-UK-LRE | Freshwater | Western Europe | 57.611 | -7.515 | 11.111 |
| PP-UK-LRE-3 | PP-UK-LRE | Freshwater | Western Europe | 57.611 | -7.515 | 11.053 |

| | | | | | | |
|---|---|---|---|---|---|---|
| PP-UK-LSC-1 | PP-UK-LSC | Freshwater | Western Europe | 57.584 | -7.236 | 8.963 |
| PP-UK-LSC-2 | PP-UK-LSC | Freshwater | Western Europe | 57.584 | -7.236 | 6.467 |
| GBR-GRO-1 | GBR-GRO | Freshwater | Western Europe | 57.615 | -7.511 | 12.118 |
| GBR-GRO-2 | GBR-GRO | Freshwater | Western Europe | 57.615 | -7.511 | 12.232 |
| GBR-GRO-3 | GBR-GRO | Freshwater | Western Europe | 57.615 | -7.511 | 12.103 |
| GBR-GRO-4 | GBR-GRO | Freshwater | Western Europe | 57.615 | -7.511 | 12.2 |
| GBR-GRO-5 | GBR-GRO | Freshwater | Western Europe | 57.615 | -7.511 | 12.422 |
| GBR-GRO-6 | GBR-GRO | Freshwater | Western Europe | 57.615 | -7.511 | 12.163 |
| PHY-CAN-TEM-01 | PP-CA-TEM | Freshwater | Western Atlantic | 47.714 | -68.917 | 8.533 |
| PHY-CAN-TEM-02 | PP-CA-TEM | Freshwater | Western Atlantic | 47.714 | -68.917 | 10.296 |
| PP-US-MAY-2 | PP-US-MAY | Freshwater | Western Atlantic | 43.343 | -70.554 | 6.983 |
| PP-US-MAY-3 | PP-US-MAY | Freshwater | Western Atlantic | 43.343 | -70.554 | 6.87 |
| PP-RU-MAG-1 | PP-RU-MAG | Freshwater | Western Pacific | 59.45 | 148.71 | 6.279 |
| PP-RU-MAG-2 | PP-RU-MAG | Freshwater | Western Pacific | 59.45 | 148.71 | 8.457 |
| PP-RU-KVA-1 | PP-RU-KVA | Freshwater | Western Pacific | 57.767 | 157.148 | 4.138 |
| PP-RU-KVA-2 | PP-RU-KVA | Freshwater | Western Pacific | 57.767 | 157.148 | 4.785 |
| PP-RU-KVA-3 | PP-RU-KVA | Freshwater | Western Pacific | 57.767 | 157.148 | 5.041 |
| PP-RU-KVA-7 | PP-RU-KVA | Freshwater | Western Pacific | 57.767 | 157.148 | 8.152 |
| PHY-RU-BLS-01 | PP-RU-BLS | Freshwater | Western Pacific | 52.757 | 156.253 | 7.69 |
| PHY-RU-BLS-02 | PP-RU-BLS | Freshwater | Western Pacific | 52.757 | 156.253 | 6.968 |
| PP-RU-GOR-6 | PP-RU-GOR | Freshwater | Western Pacific | 43.824 | 146.712 | 4.536 |
| PP-RU-GOR-7 | PP-RU-GOR | Freshwater | Western Pacific | 43.824 | 146.712 | 5.324 |
| PP-US-CRO-1 | PP-US-CRO | Freshwater | Eastern Pacific | 61.605 | -149.511 | 11.364 |
| PP-US-CRO-2 | PP-US-CRO | Freshwater | Eastern Pacific | 61.605 | -149.511 | 8.74 |
| USA-HLA-10 | US-HLA | Freshwater | Eastern Pacific | 61.591 | -149.76 | 12.477 |
| USA-HLA-11 | US-HLA | Freshwater | Eastern Pacific | 61.591 | -149.76 | 12.482 |
| USA-HLA-12 | US-HLA | Freshwater | Eastern Pacific | 61.591 | -149.76 | 12.223 |
| USA-HLA-13 | US-HLA | Freshwater | Eastern Pacific | 61.591 | -149.76 | 12.286 |
| USA-HLA-14 | US-HLA | Freshwater | Eastern Pacific | 61.591 | -149.76 | 12.293 |
| USA-HLA-15 | US-HLA | Freshwater | Eastern Pacific | 61.591 | -149.76 | 12.541 |
| USA-HLA-16 | US-HLA | Freshwater | Eastern Pacific | 61.591 | -149.76 | 12.373 |
| USA-HLA-17 | US-HLA | Freshwater | Eastern Pacific | 61.591 | -149.76 | 12.389 |
| USA-HLA-18 | US-HLA | Freshwater | Eastern Pacific | 61.591 | -149.76 | 12.406 |
| USA-HLA-19 | US-HLA | Freshwater | Eastern Pacific | 61.591 | -149.76 | 12.418 |
| USA-HLA-2 | US-HLA | Freshwater | Eastern Pacific | 61.591 | -149.76 | 12.39 |
| USA-HLA-20 | US-HLA | Freshwater | Eastern Pacific | 61.591 | -149.76 | 12.535 |
| USA-HLA-21 | US-HLA | Freshwater | Eastern Pacific | 61.591 | -149.76 | 12.49 |
| USA-HLA-3 | US-HLA | Freshwater | Eastern Pacific | 61.591 | -149.76 | 12.432 |
| USA-HLA-4 | US-HLA | Freshwater | Eastern Pacific | 61.591 | -149.76 | 12.607 |
| USA-HLA-5 | US-HLA | Freshwater | Eastern Pacific | 61.591 | -149.76 | 12.624 |
| USA-HLA-6 | US-HLA | Freshwater | Eastern Pacific | 61.591 | -149.76 | 12.543 |
| USA-HLA-7 | US-HLA | Freshwater | Eastern Pacific | 61.591 | -149.76 | 12.51 |
| USA-HLA-8 | US-HLA | Freshwater | Eastern Pacific | 61.591 | -149.76 | 12.462 |
| USA-HLA-9 | US-HLA | Freshwater | Eastern Pacific | 61.591 | -149.76 | 12.433 |
| GER-RUE-2 | GER-RUE | Marine | Baltic Sea | 54.402 | 13.21 | 13.705 |
| GER-RUE-3 | GER-RUE | Marine | Baltic Sea | 54.402 | 13.21 | 11.102 |
| GER-RUE-4 | GER-RUE | Marine | Baltic Sea | 54.402 | 13.21 | 11.011 |
| GER-RUE-5 | GER-RUE | Marine | Baltic Sea | 54.402 | 13.21 | 23.822 |
| GER-RUE-6 | GER-RUE | Marine | Baltic Sea | 54.402 | 13.21 | 23.714 |
| GER-RUE-7 | GER-RUE | Marine | Baltic Sea | 54.402 | 13.21 | 15.042 |
| 1-f | FI-HEL | Marine | Baltic Sea | 60.203 | 25.183 | 10.917 |
| 2-f | FI-HEL | Marine | Baltic Sea | 60.203 | 25.183 | 11.228 |
| 3-f | FI-HEL | Marine | Baltic Sea | 60.203 | 25.183 | 6.364 |
| 4-f | FI-HEL | Marine | Baltic Sea | 60.203 | 25.183 | 6.284 |
| 5-f | FI-HEL | Marine | Baltic Sea | 60.203 | 25.183 | 5.225 |
| 7-f | FI-HEL | Marine | Baltic Sea | 60.203 | 25.183 | 10.015 |
| SWE-BOL-60 | SWE-BOL | Marine | Baltic Sea | 63.661 | 20.212 | 9.355 |
| SWE-BOL-61 | SWE-BOL | Marine | Baltic Sea | 63.661 | 20.212 | 11.618 |
| SWE-BOL-62 | SWE-BOL | Marine | Baltic Sea | 63.661 | 20.212 | 10.719 |
| SWE-BOL-63 | SWE-BOL | Marine | Baltic Sea | 63.661 | 20.212 | 11.58 |
| SWE-BOL-64 | SWE-BOL | Marine | Baltic Sea | 63.661 | 20.212 | 12.116 |
| SWE-BOL-65 | SWE-BOL | Marine | Baltic Sea | 63.661 | 20.212 | 11.034 |
| FIN-KIV-1 | FIN-KIV | Marine | Baltic Sea | 65.008 | 25.436 | 16.484 |
| FIN-KIV-2 | FIN-KIV | Marine | Baltic Sea | 65.008 | 25.436 | 15.889 |
| FIN-KIV-3 | FIN-KIV | Marine | Baltic Sea | 65.008 | 25.436 | 15.52 |
| FIN-KIV-5 | FIN-KIV | Marine | Baltic Sea | 65.008 | 25.436 | 14.902 |
| FIN-KIV-6 | FIN-KIV | Marine | Baltic Sea | 65.008 | 25.436 | 16.813 |
| FIN-KIV-7 | FIN-KIV | Marine | Baltic Sea | 65.008 | 25.436 | 17.262 |
| RUS-LEV-10 | RUS-LEV | Marine | White and Barents Seas | 66.374 | 33.769 | 10.913 |
| RUS-LEV-11 | RUS-LEV | Marine | White and Barents Seas | 66.374 | 33.769 | 10.919 |
| RUS-LEV-3 | RUS-LEV | Marine | White and Barents Seas | 66.374 | 33.769 | 11.354 |
| RUS-LEV-7 | RUS-LEV | Marine | White and Barents Seas | 66.374 | 33.769 | 11.347 |
| RUS-LEV-8 | RUS-LEV | Marine | White and Barents Seas | 66.374 | 33.769 | 10.986 |
| RUS-LEV-9 | RUS-LEV | Marine | White and Barents Seas | 66.374 | 33.769 | 11.37 |
| DEN-NOR-10 | DEN-NOR | Marine | North Sea | 54.985 | 8.659 | 10.735 |
| DEN-NOR-12 | DEN-NOR | Marine | North Sea | 54.985 | 8.659 | 12.122 |
| DEN-NOR-13 | DEN-NOR | Marine | North Sea | 54.985 | 8.659 | 14.219 |
| DEN-NOR-6 | DEN-NOR | Marine | North Sea | 54.985 | 8.659 | 11.014 |
| DEN-NOR-7 | DEN-NOR | Marine | North Sea | 54.985 | 8.659 | 10.91 |
| DEN-NOR-9 | DEN-NOR | Marine | North Sea | 54.985 | 8.659 | 11.192 |
| DEN-NOR-14 | DEN-NOR | Marine | North Sea | 54.985 | 8.659 | 9.304 |
| DEN-NOR-15 | DEN-NOR | Marine | North Sea | 54.985 | 8.659 | 9.52 |
| DEN-NOR-16 | DEN-NOR | Marine | North Sea | 54.985 | 8.659 | 10.614 |
| SWE-FIS-35 | SWE-FIS | Marine | North Sea | 58.233 | 11.4 | 12.396 |
| SWE-FIS-36 | SWE-FIS | Marine | North Sea | 58.233 | 11.4 | 12.787 |
| SWE-FIS-40 | SWE-FIS | Marine | North Sea | 58.233 | 11.4 | 12.669 |
| SWE-FIS-42 | SWE-FIS | Marine | North Sea | 58.233 | 11.4 | 12.443 |
| SWE-FIS-43 | SWE-FIS | Marine | North Sea | 58.233 | 11.4 | 18.584 |
| SWE-FIS-45 | SWE-FIS | Marine | North Sea | 58.233 | 11.4 | 16.461 |
| SWE-FIS-49 | SWE-FIS | Marine | North Sea | 58.233 | 11.4 | 15.865 |
| SWE-FIS-52 | SWE-FIS | Marine | North Sea | 58.233 | 11.4 | 15.209 |
| SWE-FIS-53 | SWE-FIS | Marine | North Sea | 58.233 | 11.4 | 12.614 |

**Supplementary Table 3** | Populations in Figure 1b,c.

| SPECIES | POP INDEX | POP ID | ECOTYPE | REGION | LOCATION |
|---|---|---|---|---|---|
| Three-spined stickeback | POP 1 | Ran-M | Marine | Atlantic | EU |
| Three-spined stickeback | POP 2 | BS-3sp | Marine | Atlantic | EU |
| Three-spined stickeback | POP 3 | FIS | Marine | Atlantic | EU |
| Three-spined stickeback | POP 4 | KRI | Marine | Atlantic | EU |
| Three-spined stickeback | POP 5 | LEV | Marine | Atlantic | EU |
| Three-spined stickeback | POP 6 | IND | Marine | Atlantic | EU |
| Three-spined stickeback | POP 7 | SBJ | Marine | Atlantic | EU |
| Three-spined stickeback | POP 8 | BAR | Marine | Atlantic | EU |
| Three-spined stickeback | POP 9 | PRI | Marine | Atlantic | EU |
| Three-spined stickeback | POP 10 | HAL | Marine | Atlantic | Western Atlantic |
| Three-spined stickeback | POP 11 | FOR | Marine | Atlantic | Western Atlantic |
| Three-spined stickeback | POP 12 | MAI | Marine | Atlantic | Western Atlantic |
| Three-spined stickeback | POP 13 | SHI | Marine | Pacific | Western Pacific |
| Three-spined stickeback | POP 14 | ASH | Marine | Pacific | Western Pacific |
| Three-spined stickeback | POP 15 | KHA | Marine | Pacific | Western Pacific |
| Three-spined stickeback | POP 16 | ANA | Marine | Pacific | Western Pacific |
| Three-spined stickeback | POP 17 | BUT | Freshwater | Atlantic | EU |
| Three-spined stickeback | POP 18 | QUI | Freshwater | Atlantic | EU |
| Three-spined stickeback | POP 19 | KIN | Freshwater | Atlantic | EU |
| Three-spined stickeback | POP 20 | MYR | Freshwater | Atlantic | EU |
| Three-spined stickeback | POP 21 | MYV | Freshwater | Atlantic | EU |
| Three-spined stickeback | POP 22 | FAR | Freshwater | Atlantic | EU |
| Three-spined stickeback | POP 23 | TAK | Freshwater | Atlantic | EU |
| Three-spined stickeback | POP 24 | SKF | Freshwater | Atlantic | EU |
| Three-spined stickeback | POP 25 | NEV | Freshwater | Atlantic | EU |
| Three-spined stickeback | POP 26 | SLI | Freshwater | Atlantic | EU |
| Three-spined stickeback | POP 27 | VAT | Freshwater | Atlantic | EU |
| Three-spined stickeback | POP 28 | ROT | Freshwater | Atlantic | EU |
| Three-spined stickeback | POP 29 | KEV | Freshwater | Atlantic | EU |
| Three-spined stickeback | POP 30 | KVA | Freshwater | Atlantic | EU |
| Three-spined stickeback | POP 31 | MAS | Freshwater | Atlantic | EU |
| Three-spined stickeback | POP 32 | BOL | Freshwater | Atlantic | EU |
| Three-spined stickeback | POP 33 | LPD | Freshwater | Atlantic | Western Atlantic |
| Three-spined stickeback | POP 34 | PF-3sp | Freshwater | Pacific | Western Pacific |
| Three-spined stickeback | POP 35 | OTS | Freshwater | Pacific | Western Pacific |
| Three-spined stickeback | POP 36 | OIR | Freshwater | Pacific | Western Pacific |
| Three-spined stickeback | POP 37 | ALA | Freshwater | Pacific | Eastern Pacific |
| Three-spined stickeback | POP 38 | MIS | Freshwater | Pacific | Eastern Pacific |
| Three-spined stickeback | POP 39 | BEV | Freshwater | Pacific | Eastern Pacific |
| Three-spined stickeback | POP 40 | PYE | Freshwater | Pacific | Eastern Pacific |

| SPECIES | POP INDEX | POP ID | ECOTYPE | REGION | LOCATION |
|---|---|---|---|---|---|
| Nine-spined stickeback | POP 1 | SWE-FIS | Marine | Atlantic | EU |
| Nine-spined stickeback | POP 2 | DEN-NOR | Marine | Atlantic | EU |
| Nine-spined stickeback | POP 3 | RUS-LEV | Marine | Atlantic | EU |
| Nine-spined stickeback | POP 4 | FIN-KIV | Marine | Atlantic | EU |
| Nine-spined stickeback | POP 5 | SWE-BOL | Marine | Atlantic | EU |
| Nine-spined stickeback | POP 6 | FI-HEL | Marine | Atlantic | EU |
| Nine-spined stickeback | POP 7 | GER-RUE | Marine | Atlantic | EU |
| Nine-spined stickeback | POP 8 | GBR-GRO | Freshwater | Atlantic | EU |
| Nine-spined stickeback | POP 9 | PP-UK-LSC | Freshwater | Atlantic | EU |
| Nine-spined stickeback | POP 10 | PP-UK-LRE | Freshwater | Atlantic | EU |
| Nine-spined stickeback | POP 11 | PP-UK-HAR | Freshwater | Atlantic | EU |
| Nine-spined stickeback | POP 12 | BEL-MAL | Freshwater | Atlantic | EU |
| Nine-spined stickeback | POP 13 | PP-FR-MAR | Freshwater | Atlantic | EU |
| Nine-spined stickeback | POP 14 | NOR-UGE | Freshwater | Atlantic | EU |
| Nine-spined stickeback | POP 15 | PP-NO-STO | Freshwater | Atlantic | EU |
| Nine-spined stickeback | POP 16 | PP-FIN-PAL | Freshwater | Atlantic | EU |
| Nine-spined stickeback | POP 17 | DEN-RES | Freshwater | Atlantic | EU |
| Nine-spined stickeback | POP 18 | SWE-HAN | Freshwater | Atlantic | EU |
| Nine-spined stickeback | POP 19 | SWE-NAV | Freshwater | Atlantic | EU |
| Nine-spined stickeback | POP 20 | SWE-BYN | Freshwater | Atlantic | EU |
| Nine-spined stickeback | POP 21 | SWE-ABB | Freshwater | Atlantic | EU |
| Nine-spined stickeback | POP 22 | VEN-AAN | Freshwater | Atlantic | EU |
| Nine-spined stickeback | POP 23 | FIN-KEV | Freshwater | Atlantic | EU |
| Nine-spined stickeback | POP 24 | FIN-PUL | Freshwater | Atlantic | EU |
| Nine-spined stickeback | POP 25 | RUS-KRU | Freshwater | Atlantic | EU |
| Nine-spined stickeback | POP 26 | RUS-MAS | Freshwater | Atlantic | EU |
| Nine-spined stickeback | POP 27 | RUS-BOL | Freshwater | Atlantic | EU |
| Nine-spined stickeback | POP 28 | PP-US-MAY | Freshwater | Atlantic | Western Atlantic |
| Nine-spined stickeback | POP 29 | PP-CA-TEM | Freshwater | Atlantic | Western Atlantic |
| Nine-spined stickeback | POP 30 | PP-RU-GOR | Freshwater | Pacific | Western Pacific |
| Nine-spined stickeback | POP 31 | PP-RU-BLS | Freshwater | Pacific | Western Pacific |
| Nine-spined stickeback | POP 32 | PP-RU-KVA | Freshwater | Pacific | Western Pacific |
| Nine-spined stickeback | POP 33 | PP-RU-MAG | Freshwater | Pacific | Western Pacific |
| Nine-spined stickeback | POP 34 | US-HLA | Freshwater | Pacific | Eastern Pacific |
| Nine-spined stickeback | POP 35 | PP-US-CRO | Freshwater | Pacific | Eastern Pacific |

### **Supplementary Table 4** | Information on LD-clusters of three- and nine-spined sticklebacks.

**Regional EMMAX analyses of the association between “synthetic multi-locus alleles” and ecotype based on loci from significant LD-clusters for a given outlier genomic region (after corrections for p-value inflation or permutation)**

| Sp | Chr | Pos | Region | No. loci | Fst (Q95%) | C | Range | P_min | cor_A\|WB_Sea | cor_unrl\|WB_Sea | cor_A\|Baltic | cor_unrl\|Baltic | cor_A\|NorthSea | cor_unrl\|NorthSea | cor_A\|Norwegian | cor_unrl\|Norwegian | P_A\|WB_Sea | P_unrl\|WB_Sea | P_A_GC\|WB_Sea | P_unrl_GC\|WB_Sea |
|---|---|---|---|---|---|---|---|---|---|---|---|---|---|---|---|---|---|---|---|---|
| 3sp | Chr1 | 23981492 | Chr1_2 | 1341 | 0.856 | 0.958 | 650035 | 1.00E-04 | 0.243 | 0.486 | 0.39 | 0.182 | 0.78 | 0.754 | 0.544 | 0.856 | 0.552 | 0 | 0.176 | 0.007 |
| 3sp | Chr5 | 109173367 | Chr5_1 | 50 | 0.835 | 0.896 | 18569 | 1.00E-04 | 0.261 | 0.437 | 0.454 | 0.654 | 0.402 | 0.237 | 0.531 | 0.802 | 0.462 | 0.003 | 0.144 | 0.017 |
| 3sp | Chr12 | 257876940 | Chr12_1 | 77 | 0.558 | 0.701 | 1.54E+05 | 1.00E-04 | 0.148 | 0 | 0.212 | 0.257 | 0.405 | 0.437 | 0.305 | 0.638 | 0.934 | 0.999 | 0.428 | 0.96 |
| 3sp | Chr20 | 366142830 | Chr20_1 | 183 | 0.835 | 0.625 | 8.22E+04 | 1.00E-04 | 0.118 | 0.219 | 0.492 | 0.601 | 0.399 | 0.307 | 0.61 | 0.86 | 0.964 | 0.535 | 0.543 | 0.283 |
| 3sp | Chr4 | 87044465 | Chr4_2 | 135 | 0.760 | 0.583 | 4.23E+04 | 3.00E-04 | 0.055 | 0.032 | 0.373 | 0.386 | 0.577 | 0.504 | 0.613 | 0.878 | 0.978 | 0.999 | 0.792 | 0.918 |
| 3sp | Chr12 | 260204121 | Chr12_2 | 52 | 0.858 | 0.562 | 25943 | 1.00E-04 | 0.077 | 0.295 | 0.374 | 0.453 | 0.442 | 0.313 | 0.477 | 0.754 | 0.973 | 0.166 | 0.707 | 0.132 |
| 3sp | Chr7 | 149929482 | Chr7_4 | 252 | 0.342 | 0.542 | 1.39E+06 | 1.00E-04 | 0.118 | 0.276 | 0.263 | 0.335 | 0.385 | 0.363 | 0.319 | 0.658 | 0.97 | 0.235 | 0.537 | 0.164 |
| 3sp | Chr20 | 374832771 | Chr20_3 | 59 | 0.857 | 0.444 | 1.60E+04 | 1.00E-04 | 0.274 | 0.454 | 0.327 | 0.443 | 0.327 | 0.212 | 0.522 | 0.811 | 0.387 | 0 | 0.122 | 0.013 |
| 3sp | Chr13 | 282190862 | Chr13_1 | 52 | 0.363 | 0.417 | 2.33E+04 | 1.00E-04 | 0.155 | 0.352 | 0.13 | 0.361 | 0.308 | 0.493 | 0.473 | 0.792 | 0.94 | 0.055 | 0.406 | 0.066 |
| 3sp | Chr4 | 94084384 | Chr4_3 | 255 | 0.565 | 0.389 | 99349 | 3.00E-04 | 0.155 | 0.055 | 0.164 | 0.245 | 0.367 | 0.245 | 0.585 | 0.867 | 0.935 | 0.988 | 0.411 | 0.827 |
| 3sp | Chr7 | 153052832 | Chr7_5 | 97 | 0.722 | 0.361 | 3.83E+04 | 1.00E-04 | 0.084 | 0.179 | 0.161 | 0.105 | 0.318 | 0.519 | 0.465 | 0.78 | 0.996 | 0.759 | 0.683 | 0.395 |
| 3sp | Chr3 | 58199692 | Chr3_2 | 36 | 0.791 | 0.333 | 13749 | 1.00E-04 | 0.184 | 0.351 | 0.202 | 0.152 | 0.339 | 0.182 | 0.631 | 0.887 | 0.878 | 0.058 | 0.322 | 0.067 |
| 3sp | Chr4 | 94099576 | Chr4_4 | 127 | 0.531 | 0.229 | 54496 | 4.00E-04 | 0 | 0.055 | 0.219 | 0.396 | 0.439 | 0.469 | 0.354 | 0.695 | 0.954 | 0.998 | 0.971 | 0.806 |
| 3sp | Chr17 | 342328537 | Chr17_1 | 11 | 0.780 | 0.194 | 3.61E+02 | 1.00E-04 | 0.161 | 0.351 | 0.13 | 0.342 | 0.335 | 0.341 | 0.205 | 0.452 | 0.947 | 0.057 | 0.388 | 0.067 |
| 3sp | Chr20 | 382155619 | Chr20_4 | 11 | 0.443 | 0.194 | 3.40E+05 | 1.00E-04 | 0.077 | 0.1 | 0 | 0.063 | 0.182 | 0.281 | 0.19 | 0.438 | 0.991 | 0.955 | 0.691 | 0.659 |
| 3sp | Chr20 | 382155619 | Chr20_5 | 12 | 0.298 | 0.194 | 4.16E+05 | 1.00E-04 | 0.122 | 0.378 | 0.114 | 0.114 | 0.335 | 0.425 | 0.045 | 0.214 | 0.968 | 0.032 | 0.523 | 0.045 |
| 3sp | Chr7 | 157882037 | Chr7_1 | 20 | 0.334 | 0.188 | 5158 | 1.02E-02 | 0.21 | 0.377 | 0.089 | 0.167 | 0.237 | 0.361 | 0.145 | 0.327 | 0.772 | 0.03 | 0.252 | 0.046 |
| 3sp | Chr7 | 153328122 | Chr7_2 | 11 | 0.179 | 0.174 | 1587 | 4.00E-04 | 0.045 | 0 | 0.138 | 0.148 | 0.232 | 0.161 | 0.324 | 0.638 | 0.972 | 0.998 | 0.814 | 0.965 |
| 3sp | Chr7 | 153328122 | Chr7_3 | 10 | 0.152 | 0.174 | 1990 | 4.00E-04 | 0.114 | 0 | 0.13 | 0.173 | 0.187 | 0.224 | 0.277 | 0.524 | 0.955 | 0.971 | 0.547 | 0.987 |
| 3sp | Chr1 | 3324448 | Chr1_1 | 35 | 0.652 | 0.167 | 8341 | 0.0058 | 0.089 | 0.105 | 0.386 | 0.694 | 0.377 | 0.369 | 0.57 | 0.844 | 0.994 | 0.961 | 0.66 | 0.647 |
| 3sp | Chr9 | 208559564 | Chr9_2 | 59 | 0.278 | 0.111 | 150251 | 1.40E-02 | 0.126 | 0.31 | 0.055 | 0.032 | 0.2 | 0.148 | 0.114 | 0.313 | 0.964 | 0.124 | 0.502 | 0.113 |
| 3sp | Chr20 | 366173761 | Chr20_2 | 16 | 0.808 | 0.097 | 5684 | 9.00E-04 | 0.077 | 0.148 | 0.122 | 0.055 | 0.497 | 0.626 | 0.504 | 0.802 | 0.989 | 0.846 | 0.702 | 0.494 |
| 3sp | Chr3 | 63993066 | Chr3_1 | 16 | 0.128 | 0.09 | 8367 | 2.10E-03 | 0.207 | 0.338 | 0 | 0.195 | 0.283 | 0.316 | 0.141 | 0.351 | 0.762 | 0.071 | 0.256 | 0.079 |
| 3sp | Chr18 | 353174923 | Chr18_1 | 12 | 0.842 | 0.09 | 3383 | 4.10E-03 | 0.333 | 0.439 | 0 | 0.173 | 0.122 | 0.161 | 0.436 | 0.729 | 0.107 | 0.002 | 0.053 | 0.016 |
| 3sp | Chr4 | 96188022 | Chr4_1 | 12 | 0.086 | 0.083 | 692 | 2.05E-02 | 0 | 0.148 | 0.409 | 0.529 | 0.283 | 0.307 | 0.377 | 0.691 | 0.993 | 0.868 | 0.946 | 0.489 |
| 3sp | Chr9 | 202542085 | Chr9_1 | 55 | 0.527 | 0.056 | 469761 | 0.024 | 0.173 | 0.134 | 0.302 | 0.394 | 0.379 | 0.432 | 0.318 | 0.592 | 0.906 | 0.886 | 0.351 | 0.541 |
| 9sp | Chr20 | 451628138 | Chr20_3 | 55 | 0.089 | 0.444 | 19197 | 1.00E-04 | 0.736 | 0.782 | 0.464 | 0.622 | 0.1 | 0.032 | 0.435 | 0.145 | 0 | 0 | 0 | 0 |
| 9sp | Chr4 | 109387633 | Chr4_1 | 27 | 0.498 | 0.417 | 5519 | 1.00E-04 | 0.366 | 0.486 | 0.315 | 0.489 | 0.122 | 0.152 | 0.063 | 0.285 | 0.009 | 0 | 0.004 | 0.001 |
| 9sp | Chr18 | 395906216 | Chr18_2 | 33 | 0.008 | 0.333 | 2856 | 1.00E-04 | 0.447 | 0.601 | 0.551 | 0.503 | 0.404 | 0.21 | 0.375 | 0.182 | 0 | 0 | 0 | 0 |
| 9sp | Chr20 | 436609240 | Chr20_1 | 58 | 0.885 | 0.299 | 10410 | 4.00E-04 | 0.414 | 0.335 | 0.412 | 0.373 | 0.155 | 0.095 | 0.118 | 0.2 | 0 | 0.022 | 0.001 | 0.029 |
| 9sp | Chr18 | 408488264 | Chr18_1 | 109 | 0.196 | 0.194 | 153576 | 8.00E-04 | 0.387 | 0.232 | 0.032 | 0 | 0.195 | 0.406 | 0.161 | 0.161 | 0.014 | 0.012 | 0.002 | 0.145 |
| 9sp | Chr14 | 311800447 | Chr14_1 | 54 | 0.031 | 0.111 | 5783 | 1.00E-04 | 0.055 | 0.268 | 0.195 | 0.436 | 0.126 | 0.184 | 0.161 | 0.055 | 0.567 | 0.022 | 0.702 | 0.089 |
| 9sp | Chr13 | 288161750 | Chr13_1 | 19 | 0.268 | 0.104 | 169638 | 1.36E-02 | 0.349 | 0.424 | 0.578 | 0.74 | 0.055 | 0 | 0.114 | 0.285 | 0.009 | 0 | 0.006 | 0.004 |
| 9sp | Chr3 | 80648276 | Chr3_1 | 10 | 0.594 | 0.083 | 124 | 8.50E-03 | 0.369 | 0.399 | 0.212 | 0.31 | 0.13 | 0.11 | 0.263 | 0.095 | 0.009 | 0 | 0.003 | 0.007 |
| 9sp | Chr20 | 448139269 | Chr20_2 | 14 | 0.572 | 0.083 | 104037 | 8.50E-03 | 0.385 | 0.391 | 0.114 | 0.184 | 0.197 | 0.182 | 0.321 | 0.045 | 0.014 | 0 | 0.002 | 0.009 |
| 9sp | Chr18 | 408488264 | Chr18_1 | 109 | 0.196 | 0.15 | 153576 | 5.00E-04 | 0.387 | 0.232 | 0.032 | 0 | 0.195 | 0.406 | 0.161 | 0.161 | 0.016 | 0.009 | 0.002 | 0.132 |
| 9sp | Chr14 | 311800447 | Chr14_2 | 54 | 0.031 | 0.133 | 5783 | 1.00E-04 | 0.055 | 0.268 | 0.195 | 0.436 | 0.126 | 0.184 | 0.161 | 0.055 | 0.549 | 0.026 | 0.699 | 0.079 |
| 9sp | Chr13 | 288161750 | Chr13_1 | 19 | 0.268 | 0.111 | 169638 | 0.0146 | 0.349 | 0.424 | 0.578 | 0.74 | 0.055 | 0 | 0.114 | 0.285 | 0.008 | <0.001 | 0.005 | 0.003 |
| 9sp | Chr3 | 80648276 | Chr3_1 | 10 | 0.594 | 0.083 | 124 | 0.0091 | 0.369 | 0.399 | 0.212 | 0.31 | 0.13 | 0.11 | 0.263 | 0.095 | 0.008 | <0.001 | 0.003 | 0.006 |
| 9sp | Chr20 | 448139269 | Chr20_1 | 14 | 0.572 | 0.083 | 104037 | 0.0091 | 0.385 | 0.391 | 0.114 | 0.184 | 0.197 | 0.182 | 0.321 | 0.045 | <0.001 | <0.001 | 0.002 | 0.007 |

| P_A\|Baltic | P_unrl\|Baltic | P_A_GC\|Baltic | P_unrl_GC\|Baltic | P_A\|NorthSea | P_unrl\|NorthSea | P_A_GC\|NorthSea | P_unrl_GC\|NorthSea | P_A\|Norwegian | P_unrl\|Norwegian | P_A_GC\|Norwegian | P_unrl_GC\|Norwegian |
|---|---|---|---|---|---|---|---|---|---|---|---|
| 0.077 | 0.857 | 0.041 | 0.451 | 0 | 0 | 0 | 0 | 0.001 | 0 | 0.002 | 0 |
| 0.01 | 0 | 0.015 | 0 | 0.031 | 0.418 | 0.03 | 0.293 | 0.001 | 0 | 0.002 | 0 |
| 0.817 | 0.544 | 0.304 | 0.258 | 0.031 | 0.017 | 0.029 | 0.032 | 0.19 | 0 | 0.111 | 0 |
| 0.003 | 0 | 0.007 | 0.002 | 0.036 | 0.176 | 0.032 | 0.159 | 0 | 0 | 0 | 0 |
| 0.097 | 0.065 | 0.053 | 0.071 | 0 | 0.002 | 0.001 | 0.011 | 0 | 0 | 0 | 0 |
| 0.1 | 0.015 | 0.051 | 0.028 | 0.015 | 0.171 | 0.016 | 0.148 | 0.004 | 0 | 0.007 | 0 |
| 0.555 | 0.162 | 0.194 | 0.127 | 0.057 | 0.077 | 0.04 | 0.085 | 0.164 | 0 | 0.094 | 0 |
| 0.228 | 0.018 | 0.095 | 0.033 | 0.146 | 0.413 | 0.088 | 0.358 | 0.001 | 0 | 0.003 | 0 |
| 0.974 | 0.107 | 0.547 | 0.096 | 0.193 | 0.004 | 0.11 | 0.013 | 0.004 | 0 | 0.008 | 0 |
| 0.949 | 0.596 | 0.435 | 0.286 | 0.08 | 0.414 | 0.051 | 0.275 | 0.001 | 0 | 0.001 | 0 |
| 0.941 | 0.908 | 0.447 | 0.682 | 0.172 | 0.002 | 0.099 | 0.008 | 0.004 | 0 | 0.01 | 0 |
| 0.853 | 0.873 | 0.33 | 0.537 | 0.14 | 0.498 | 0.075 | 0.439 | 0 | 0 | 0 | 0 |
| 0.801 | 0.051 | 0.286 | 0.062 | 0.015 | 0.008 | 0.016 | 0.019 | 0.08 | 0 | 0.059 | 0 |
| 0.961 | 0.148 | 0.552 | 0.117 | 0.139 | 0.121 | 0.08 | 0.111 | 0.572 | 0.005 | 0.305 | 0.024 |
| 0.997 | 0.96 | 0.926 | 0.814 | 0.337 | 0.251 | 0.374 | 0.202 | 0.553 | 0.005 | 0.349 | 0.03 |
| 0.952 | 0.927 | 0.608 | 0.65 | 0.132 | 0.017 | 0.08 | 0.038 | 0.782 | 0.132 | 0.846 | 0.347 |
| 0.968 | 0.84 | 0.682 | 0.487 | 0.384 | 0.081 | 0.237 | 0.088 | 0.721 | 0.035 | 0.484 | 0.123 |
| 0.979 | 0.829 | 0.519 | 0.544 | 0.35 | 0.432 | 0.247 | 0.5 | 0.161 | 0 | 0.088 | 0 |
| 0.987 | 0.864 | 0.547 | 0.468 | 0.409 | 0.43 | 0.362 | 0.325 | 0.273 | 0 | 0.151 | 0.007 |
| 0.078 | 0 | 0.043 | 0 | 0.07 | 0.072 | 0.045 | 0.08 | 0.001 | 0 | 0.001 | 0 |
| 0.988 | 0.821 | 0.817 | 0.91 | 0.409 | 0.309 | 0.328 | 0.537 | 0.667 | 0.044 | 0.592 | 0.144 |
| 0.949 | 0.907 | 0.571 | 0.831 | 0.003 | 0 | 0.005 | 0.001 | 0.001 | 0 | 0.004 | 0 |
| 0.97 | 0.82 | 0.978 | 0.408 | 0.254 | 0.172 | 0.147 | 0.143 | 0.668 | 0.023 | 0.497 | 0.095 |
| 0.983 | 0.841 | 0.927 | 0.472 | 0.376 | 0.539 | 0.569 | 0.496 | 0.014 | 0 | 0.016 | 0 |
| 0.049 | 0.001 | 0.031 | 0.008 | 0.236 | 0.184 | 0.149 | 0.157 | 0.05 | 0 | 0.042 | 0 |
| 0.349 | 0.05 | 0.128 | 0.064 | 0.065 | 0.017 | 0.043 | 0.034 | 0.161 | 0 | 0.096 | 0.001 |
| 0.03 | 0 | 0.002 | 0 | 0.259 | 0.972 | 0.514 | 0.859 | 0.031 | 0.236 | 0.001 | 0.446 |
| 0.108 | 0.028 | 0.04 | 0.005 | 0.246 | 0.404 | 0.422 | 0.417 | 0.288 | 0.169 | 0.679 | 0.102 |
| 0 | 0.028 | 0 | 0.003 | 0.037 | 0.256 | 0.003 | 0.247 | 0.063 | 0.245 | 0.007 | 0.322 |
| 0.03 | 0.028 | 0.006 | 0.041 | 0.434 | 0.316 | 0.297 | 0.628 | 0.289 | 0.233 | 0.428 | 0.27 |
| 0.865 | 1 | 0.881 | 0.947 | 0.267 | 0.031 | 0.185 | 0.014 | 0.279 | 0.378 | 0.277 | 0.383 |
| 0.241 | 0.028 | 0.22 | 0.014 | 0.286 | 0.307 | 0.401 | 0.318 | 0.252 | 0.704 | 0.281 | 0.792 |
| 0 | 0 | 0 | 0 | 0.512 | 1 | 0.713 | 0.998 | 0.202 | 0.181 | 0.457 | 0.101 |
| 0.279 | 0.121 | 0.182 | 0.099 | 0.402 | 0.285 | 0.377 | 0.568 | 0.137 | 0.368 | 0.067 | 0.628 |
| 0.433 | 0.13 | 0.48 | 0.358 | 0.267 | 0.313 | 0.176 | 0.325 | 0.092 | 0.738 | 0.023 | 0.845 |
| 0.852 | 1 | 0.879 | 0.945 | 0.269 | 0.025 | 0.181 | 0.011 | 0.314 | 0.237 | 0.272 | 0.366 |
| 0.237 | 0.033 | 0.216 | 0.011 | 0.277 | 0.339 | 0.396 | 0.301 | 0.246 | 0.687 | 0.276 | 0.784 |
| <0.001 | <0.001 | <0.001 | <0.001 | 0.524 | 1 | 0.71 | 0.998 | 0.2 | 0.163 | 0.452 | 0.091 |
| 0.295 | 0.122 | 0.178 | 0.088 | 0.425 | 0.317 | 0.372 | 0.553 | 0.128 | 0.36 | 0.065 | 0.614 |
| 0.439 | 0.126 | 0.476 | 0.341 | 0.269 | 0.35 | 0.172 | 0.308 | 0.094 | 0.725 | 0.021 | 0.838 |

Sp: Species (3sp=three-spined sticlebacks and 9sp=nine-spined sticlebacks)

Chr: Chromosome

Region: matches outlier region in Figure 5 & 6, main text

Fst (Q95%) = 95% quantile among loci that mapped to the outlier regions from significant LD-clusters

Range=size of outlier genomic region in Bps

P_min=the lowest p-value among LD-clusters that mapped to this region

**Subsequent column names should be interpreted as follows:**

cor: correlation coefficient

P: P-value from EMMAX

A: relatedness was accounted for

unrl: individuals assumed unrelated

GC: genomic control was used to account for residual p-value inflation (otherwise permutation was used to assess significance)

Note that lineage was always included as co-factor for nine-spined sticklebacks

### Outlier population (RUS-LEV) in the IBD analyses

The current marine pathway connecting White Sea and Baltic Sea through Barents and North Seas is unlikely to reflect the real geographic distance contributing to the close phylogenetic affinity between White Sea (RUS-LEV) and Baltic Sea populations sticklebacks in our data. This because the Baltic Sea populations are believed to be colonized from the White Sea after the last glaciation (see results, Fig. 3; Feng et al. 2020). In fact, the significant decrease in IBD when incorporating the White Sea RUS-LEV into the analysis (see below) is likely to be an artefact attributable to demographic history. In fact, the close relationship between the populations from the Baltic Sea and White Sea was likely a result of the southward colonisation during the retreat of the Scandinavian ice sheet in the late Pleistocene and early Holocene (Feng et al. 2020; Glückert 1995; Guo et al. 2019). Based on this assumption, the results of IBD analyses were discussed based on the dataset excluding the outlier population RUS-LEV.

When incorporating White Sea population (RUS-LEV) of the nine-spined stickleback into the IBD analyses, the IBD of marine nine-spined sticklebacks decreased by a factor of 3.1 (MLPE, $\beta$ = 1.2e-4 *vs*. 4.0e-5; Supplementary Information Fig. 13). However, the IBD of nine-spined sticklebacks was still 7.7 times stronger than that of three-spined sticklebacks in marine habitat (MLPE, $\beta$ = 4.0e-5 *vs.* 5.1e-6; Supplementary Information Fig. 1).

**Supplementary Information Figure 1 | Isolation-by-distance (IBD) in three- and nine-spined stickleback populations.** This dataset includes the nine-spined stickleback White Sea population (RUS-LEV). IBD was tested across (a) marine and (b) freshwater three- (red) and nine-spined stickleback (blue) populations using maximum-likelihood population-effects (MLPE) model. The results of the regression coefficient (β) suggest stronger IBD in nine- than in three-spined sticklebacks: 7.7 times in the marine habitat and 23.6 times in the freshwater habitat (see results). IBD comparisons were restricted to European populations (see methods).

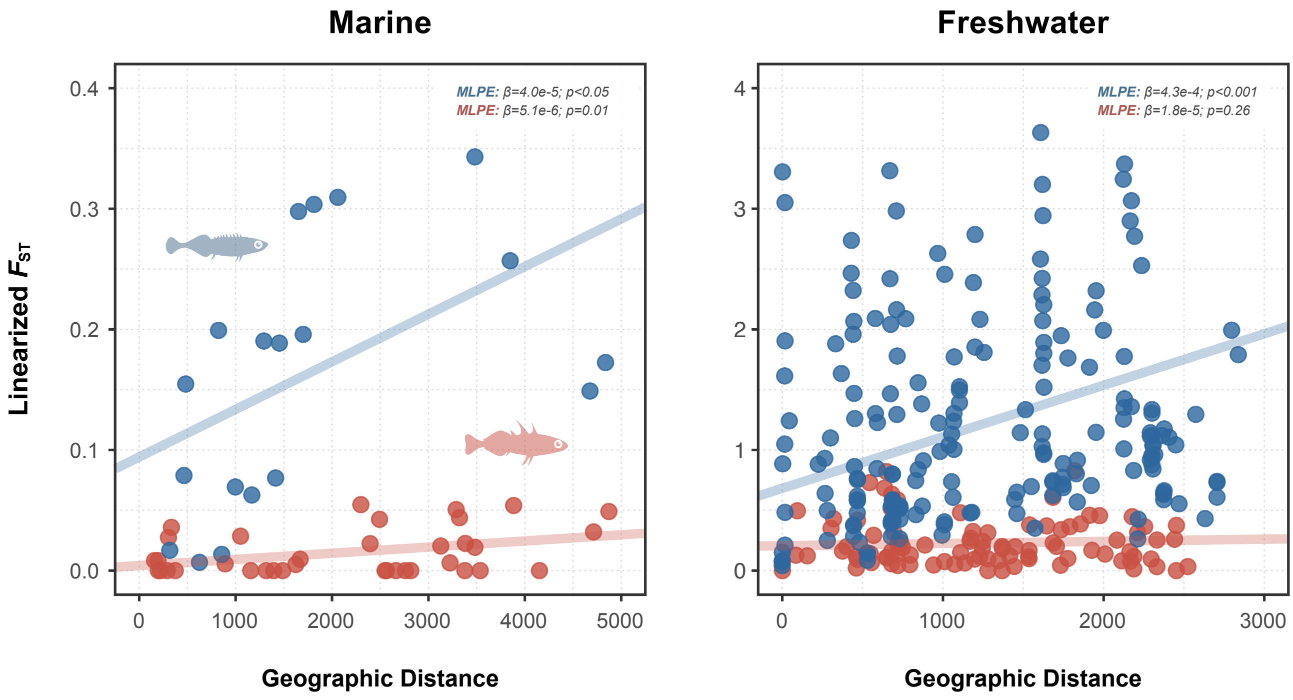

**Supplementary Information 2 | Methodological considerations**

It is generally believed that strong genetic drift makes it difficult to detect signatures of parallel evolution (Galloway et al. 2020; Hoban et al. 2016; Matthey□Doret & Whitlock 2019) and therefore the reported differences between three- and nine-spined sticklebacks in parallel evolution could be argued to be an artefact of the inherent difficulty of detecting outlier loci among highly differentiated populations. However, both LDna-complexity reduction as well as using EMMAX to test for associations between SMLAs and ecotype are likely to mitigate the many shortcomings that otherwise characterise e.g. FST-based genome scan approaches (e.g. Bierne et al. 2011; Excoffier et al. 2009; Pritchard & Di Rienzo 2010). In our analyses, EMMAX produced test statistics that were highly correlated with both $F_{ST}$ and CSS (we know already that CSS is correlated with $F_{ST}$; Jones et al. 2012), but importantly only when relatedness among individuals or other co-factors were not explicitly accounted for. *P*-value inflation in our analyses was high (up to λ~2), which was strongly reduced when relatedness was included as a random effect. However, both accounting for relatedness and correcting for p-value inflation (using GC) reduced the power to detect significant associations considerably, such that it was futile to find any outlier genomic regions without also using LDna-complexity reduction which allowed much fewer conservative corrections for multiple testing. Estimating and correcting for p-value inflation caused by relatedness is unfortunately not common practice in evolutionary genetic studies. However, nor is complexity reduction. It is thus possible that the increased power from p-value inflation roughly have counterbalanced the reduced power from unnecessarily conservative corrections for multiplicity in many previous studies. However, without re-analyses of these data sets, it is impossible to know to what extent previously detected outlier regions could in fact be artefacts of relatedness and p-value inflation.

We know that population structuring increases background differentiation in statistics estimating genetic differentiation between populations (Galloway et al. 2020; Hoban et al. 2016; Matthey□Doret & Whitlock 2019) and can also increase *p*-value inflation in genome wide association (GWA)-analyses when phenotypes are confounded by relatedness (Kang et al. 2010; Kang et al. 2008). There is thus reason to believe that these two observations are not independent, i.e. whenever background differentiation is high, we can also expect p-value inflation due to relatedness. Importantly, however, accounting for relatedness can also increase power in GWA-studies (Kang et al. 2010; Kang et al. 2008). e.g. when multiple divergent populations segregate for the same causal variants for a given phenotype. This can also be expected when multiple populations in similar habitats display high frequencies of the same genetic variants (i.e. parallel evolution); the more divergent the populations are, the stronger will the contrast be between the neutral genetic background (reflecting relatedness) and the genomic regions under selection. For the outlier regions detected here, the overall effect sizes in three-spined sticklebacks differed when including relatedness as a random effect (0.38) compared to when relatedness was not accounted for (0.28) but remained roughly the same for nine-spined sticklebacks (0.29 *vs.* 0.27, respectively). Since p-value inflation (that reflect biases due to relatedness) in three-spined sticklebacks was higher (λ=1.95) compared to nine-spined sticklebacks (λ=1.73; when including lineage as co-factor), there is no reason to suspect that our methods lacked power to detect outlier regions in nine-spined sticklebacks across comparable levels of effect sizes in both species. Despite the fact that marine and freshwater individuals were pooled across all geographic regions in our analyses, no outlier region showed high effect sizes across all geographic regions. This demonstrates that, as long as effect sizes are sufficiently high overall, our methods also have sufficient power to detect regional parallelism (in both species). This is no different from detecting significant genotype-phenotype associations in GWA-studies for phenotypes that only segregate in only one or few of the populations included in the study.

While *p*-values are important, they tell very little about effect sizes which are much more likely to reflect the extent to which a given genomic region is affected by parallel evolution. While 26 and nine outlier genomic regions were significant in three-and nine-spined sticklebacks respectively, they were to different degrees sensitive to parameter settings and varied considerably in their effect sizes. These are all important metrics to consider when determining how significant a given genomic region harboring freshwater adapted alleles are from an evolutionary/ecological perspective. However, while we can safely conclude that the marine-freshwater differentiated regions in three-spined sticklebacks outnumber such regions in nine-spined sticklebacks across a wide range of parameter/threshold values and correction methods, more detailed assessments of these outlier regions are outside the scope of this study.
